# Identification and characterization of human CD34^+^ and CD34^dim/-^ neutrophil-committed progenitors

**DOI:** 10.1101/2021.04.30.442138

**Authors:** Federica Calzetti, Giulia Finotti, Nicola Tamassia, Francisco Bianchetto-Aguilera, Monica Castellucci, Chiara Cavallini, Alessandro Mattè, Sara Gasperini, Fabio Benedetti, Massimiliano Bonifacio, Cristina Tecchio, Patrizia Scapini, Marco A. Cassatella

## Abstract

We report the identification of human CD66b^−^CD64^dim^CD115^−^ neutrophil-committed progenitors within SSC^low^CD45^dim^CD34^+^ and CD34^dim/−^ bone marrow cells, that we named neutrophil myeloblast (NMs). CD34^+^ and CD34^dim/−^NMs resulted as either CD45RA^+^ or CD45RA^−^, with CD34^+^CD45RA^−^NMs found as selectively expanded in chronic-phase chronic myeloid leukemia patients. By scRNA-seq experiments, CD34^+^ and CD34^dim/−^NMs were found to consist of combinations of four cell clusters, characterized by different maturation stages and distributed along two differentiation routes. Cell clusters were identified by neutrophil-specific gene profiles, one of them associated to an interferon-stimulated gene (ISG) signature, hence supporting recently identified expansions of mature neutrophil subsets expressing ISGs in blood of diseased individuals. Altogether, our data shed light on the very early phases of neutrophil ontogeny.

## INTRODUCTION

Recent studies, performed by single-cell sorting, transcriptional analysis, single-cell transplants and clonal tracking analysis ^1-9^, have challenged the classical hierarchical tree-like model of hematopoiesis ^10, 11^. Accordingly, hematopoietic stem cells (HSCs) are currently considered as a heterogeneous cellular population in terms of lineage potential and transcriptional profile, instead of multi-potent, homogeneous clusters of cells ^12^. In fact, it has been ascertained that hematopoiesis occurs as a continuous process along developmental trajectories following early erythroid/megakaryocyte/eosinophil/basophil and lympho/myeloid, ultimately generating mature, terminally differentiated cells. Moreover, transcriptomic profiling demonstrates that intermediate compartments of progenitors, such as common myeloid progenitors (CMPs) or granulocyte-macrophage progenitors (GMPs) ^10^, are largely composed of clusters of cells displaying lineage-selective commitment. However, even though scRNA-seq experiments are necessary to reveal their heterogeneity ^13^, the various cellular subpopulations are still identified by flow-sorting and immunophenotyping approaches.

In this scenario, conventional GMPs (cGMPs) represent the most restricted pool of myeloid progenitors in humans ^14, 15^. In fact, based on flow cytometry approaches, cGMPs have been fractionated into three main compartments ^16^, namely: i) granulocyte-monocyte-DC progenitors (GMDPs), that generate CD66b^+^ granulocytes, CD14^+^ monocytes and the three dendritic cell (DC) subsets (DC1s, DC2s and pDCs); ii) monocyte-DC progenitors (MDPs), that generate DCs and CD14^+^ monocytes; iii) common dendritic progenitors (CDPs), that are exclusively committed to DCs. By similar methodologies, but using a different panel of markers including CD64, common monocyte progenitors (cMoPs), exclusively differentiating into pre-monocytes and ultimately into CD14^+^ monocytes, were subsequently identified within cGMPs, while the existence of revised GMPs (rGMPs), generating only CD66b^+^ granulocytes and CD14^+^ monocytes, was also postulated ^17^. In this context, however, little progress has been made to characterize the ontogeny of human neutrophils, even though neutrophil - but not basophil or eosinophil - progenitors, are included in cGMPs ^3, 4, 6, 18^-^20^. For instance, a CD34^+^CD115^−^CD64^+^ fraction able to originate granulocytes was identified within the fetal bone marrow (BM) in 1996 ^21^, without, however, discriminating their eventual eosinophil, basophil or neutrophil nature. Similarly, the phenotypic and/or morphological features of the CD66b^+^ granulocytes originated by GMDPs ^16^ and rGMPs ^17^ were not exhaustively pursued. More recently, various human neutrophil progenitors have been described, precisely preNeus ^22^, hNePs ^23^, proNeus^24^ and eNePs ^25^. However, all of them express the CD66b and CD15 lineage markers, and thus resemble the promyelocyte (PM), or more mature neutrophil, stage. By contrast, in mice, CD34^+^Ly6C^+^CD115^−^ proNeu1s have been found within GMPs ^24^, thus representing the earliest identifiable progenitors along the neutrophil maturation trajectory currently identified.

Based on these premises, herein we report the identification, phenotypic characterization and single cell transcriptomic analysis of previously undescribed, human uni-lineage CD34^+^ and CD34^dim/−^ neutrophil progenitors, which we named as neutrophil myeloblast (NMs).

## RESULTS

### Characterization of myeloid progenitors within cGMPs

To identify progenitors of human neutrophils at earlier stages than PMs, we focused on SSC^low^CD45^dim^ cells present in the low-density cells of BM (BM-LDCs) (panel **III** of **Figure 1A**, and panel **V** of **Figure S1A)**. SSC^low^CD45^dim^ cells are lineage negative (**Figure S1B**) and include CD34^+^CD38^−^ HSCs, other than CD34^+^ and CD34^dim/−^ myeloid/lymphoid progenitors ^26, 27^. Moreover, although CD34^dim/−^ cells represent neglected, but important, transitional progenitor stages ^4^, they have been rarely investigated for a potential ability to generate neutrophils. Therefore, we stained BM-LDCs by a flow-cytometry antibody panel comprising key markers conventionally used to detect either BM or cord blood (CB) CD34^+^ myeloid and lymphoid progenitors, namely CD34, CD38, CD10, CD123, CD45RA, CD64 and CD115 ^14, 15, 21, 28^. By doing so, we could identify a lineage negative, SSC^low^CD45^dim^CD10^−^CD38^+^ region displaying variable CD34 and CD45RA levels (panel **V** of **Figure 1A**), and including CD34^+^CD45RA^−^, CD34^+^CD45RA^+^, CD34^dim/−^CD45RA^+^ and CD34^dim/−^CD45RA^−^ cell populations. It is noteworthy that CD34^+^CD45RA^+^ cells were also found CD135^+^ (**data not shown**), and thus represent cGMPs (panel **V** of **Figure 1A**) ^15^, while CD34^+^CD45RA^−^ cells (panel **V** of **Figure 1A**) include more immature progenitors, such as CMPs and MEPs ^14, 15^. We then focused on the cGMP region and subdivided it into a total of five discrete cell populations based on their CD123, CD115 and CD64 expression. Precisely: i) CDPs (panel **VI** of **Figure 1A**), which are strictly CD64^−^ (**data not shown**); ii) CD64^+^CD115^+^ populations, resembling the monocyte-committed progenitor population (CFU-M) ^21^, as well as cMoPs ^17^, that we provisionally named cMoP-like cells (panel **VII** of **Figure 1A**); iii) CD64^−^CD115^+^ populations resembling MDPs ^16^, that we provisionally named MDP-like cells (panel **VII** of **Figure 1A**); (iv) CD64^−^CD115^−^ populations resembling the conventional GMDPs ^16^, that we provisionally named CD64^−^GMDP-like cells (panel **VII** of **Figure 1A**); and v) CD64^dim^CD115^−^ cells, resembling previously described granulocyte progenitors ^21^(panel **VII** of **Figure 1A**).

**Figure 1.**
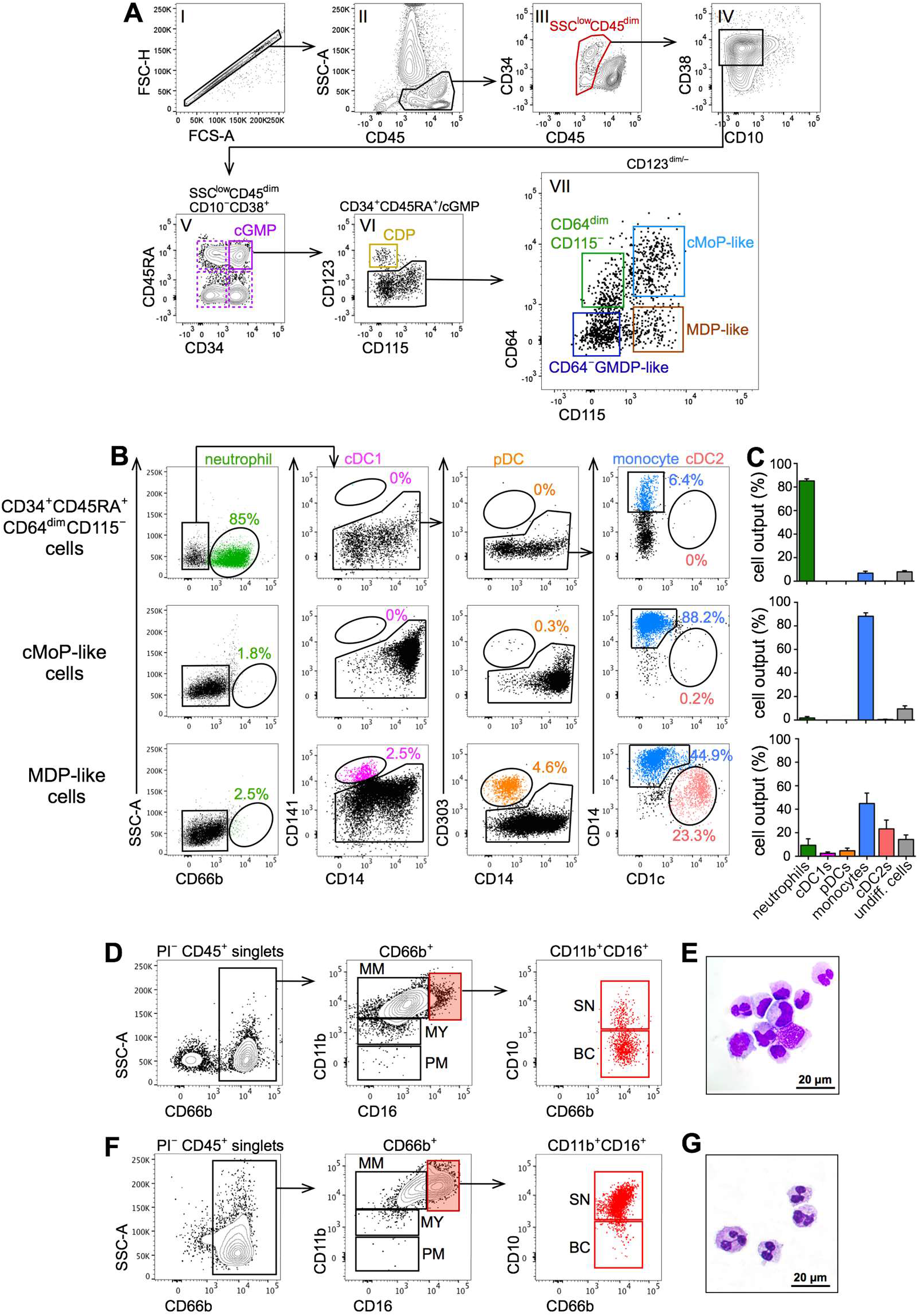
Identification of neutrophil progenitors within cGMPs. (A) Representative flow cytometry plots displaying the gating strategy to identify progenitors within the lineage negative SSC^low^CD45^dim^ region of BM-LDCs (as detailed in Figure S1, n=20). cGMPs are identified as singlet, SSC^low^CD45^dim^CD10^−^CD38^+^CD34^+^CD45RA^+^ cells, then sub-gated into five populations based on their CD123, CD64 and CD115 expression, as described in the text. (B) Representative flow cytometry plots showing the phenotypes of the cells generated from CD34^+^CD45RA^+^CD64^dim^CD115^−^, cMoP-like and MDP-like cells incubated with SFGc for 7 days (n=6-15). Values in each panel indicate the percentage of gated cells relative to live, CD45^+^ cells. (C) Bar graphs reporting the percentages (mean±SEM relative to live CD45^+^ cells, n=6-15) of the mature leukocyte types (listed in the *x* axis) generated by the experiments shown in panel B. (D,F) Representative plots displaying the phenotype of neutrophils derived from CD34^+^CD45RA^+^CD64^dim^CD115^−^ cells cultured for 7 (panel D, n=15) and 14 (panel F, n=3) days with SFGc. Neutrophils were gated as live, CD45^+^CD66b^+^ cells. (E,G) Representative morphology of neutrophils derived from CD34^+^CD45RA^+^CD64^dim^CD115^−^ cells after 7 (panel E) and 14 (panel G) days of culture (n=3).

### CD34^+^CD45RA^+^CD64^dim^CD115^−^cells are neutrophil-restricted progenitors

Recent studies reported that neutrophils, but not basophils/eosinophils, derive from cGMPs ^4, 18^-^20^. Therefore, to investigate whether CD34^+^CD45RA^+^CD64^dim^CD115^−^ cells represent neutrophil-restricted progenitors, we seeded them on top of MS-5 cells ^29^ for seven days, in a medium containing a cocktail of SCF, Flt3L and G-CSF (SFGc), and, in turn, found them to give origin mostly to CD66b^+^ cells, and poorly to monocytes (**Figure 1B** and **1C**, top panels). By contrast, SFGc-treated cMoP-like cells were found to differentiate mainly into CD14^+^ monocytes, and minimally to CD66b^+^ cells and DCs (**Figure 1B** and **1C**, middle panels), confirming their commitment to monocytes ^17^. Conversely, SFGc-treated MDP-like cells were found to originate CD141^+^CD14^−^ cDC1, CD303^+^CD14^−^ pDCs, CD14^dim/−^CD1c^+^ cDC2 and CD14^+^ monocytes (**Figure 1B** and **1C**, bottom panels), consistent with their predicted heterogeneous composition ^16, 30^. Moreover, SFGc was found to fully maintain the viability of progenitor-generated cells after 7 days (**Figure S2A**), as well as to potently promote the expansion of CD34^+^CD45RA^+^CD64^dim^CD115^−^ cells compared to cMoP-like and MDP-like cells (**Figure S2B**), consistent with its high specificity for the neutrophil lineage, and with the highest expression of CD114/G-CSFR in CD34^+^CD45RA^+^CD64^dim^CD115^−^ cells (**Figure S2C**). Phenotypic (**Figure 1D**) and morphologic (**Figure 1E**) analysis of CD66b^+^ cells derived from SFGc-treated CD34^+^CD45RA^+^CD64^dim^CD115^−^ cells revealed that they mostly consist of immature neutrophils, being composed of: CD11b^−^CD16^−^ PMs; CD11b^dim/+^CD16^−^ myelocytes (MYs); CD11b^+^CD16^+^ metamyelocytes (MMs); CD11b^+^CD16^++^CD10^−^ band cells (BCs); and CD11b^+^CD16^++^CD10^+^ segmented neutrophils (SNs) (for the respective percentages, see also second row of **Figure 2D**). Consistently, CD34^+^CD45RA^+^CD64^dim^CD115^−^ cells incubated with SFGc for 14 days originated more mature stages of neutrophils, such as BCs and SNs (**Figure 1F**), as also confirmed by the morphology of their nuclei (**Figure 1G**). Moreover, specificity of lineage commitment to neutrophils was demonstrated by culturing CD34^+^CD45RA^+^CD64^dim^CD115^−^ cells with SCF, Flt3L and GM-CSF (SFG) (**Figure S2D)**, a condition that induced monocytes or monocytes and DCs by, respectively, cMoP-like and MDP-like cells (**Figure S2E** and **F)**, as reported for cMoPs and MDPs ^17, 29^.

**Figure 2.**
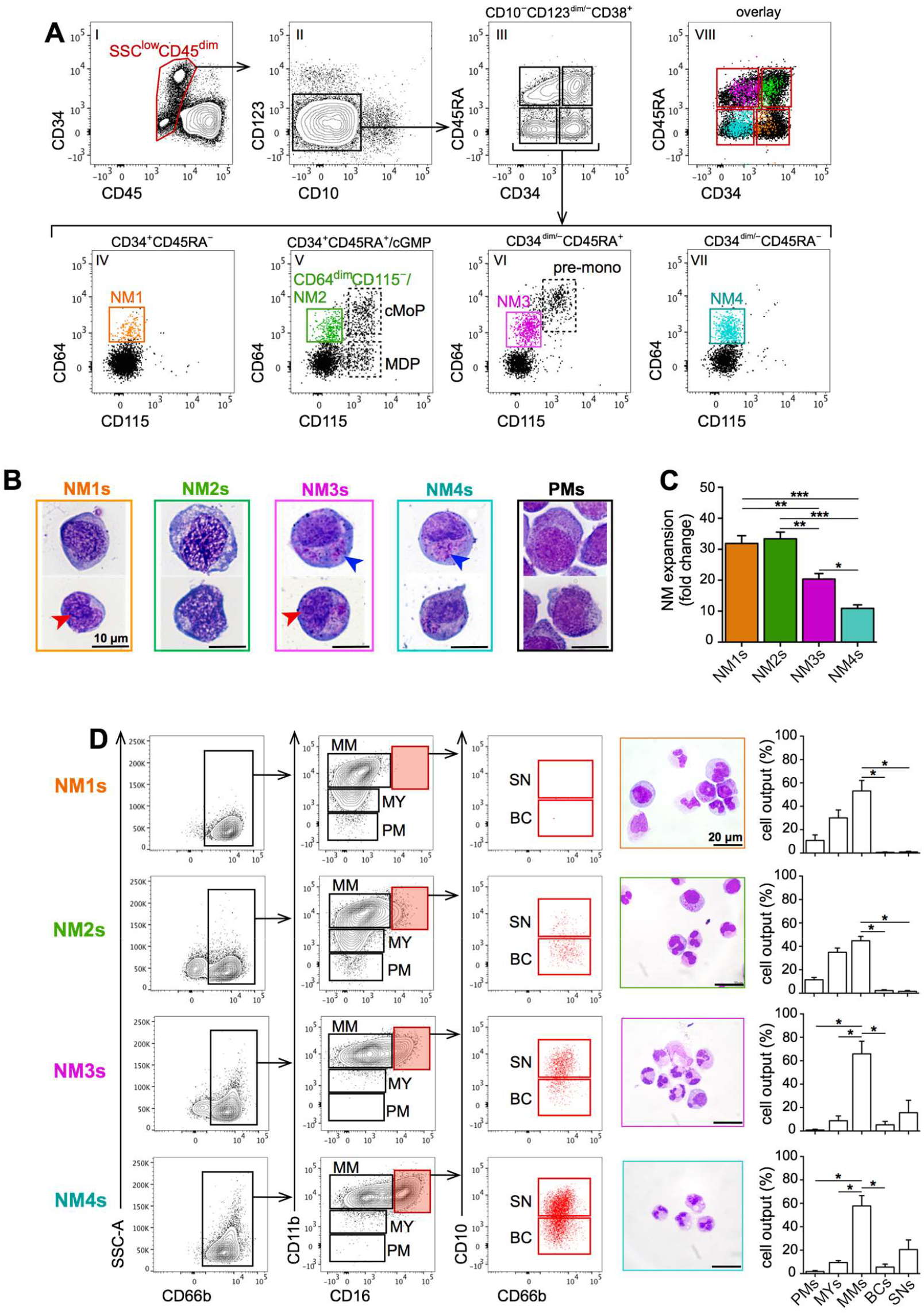
Identification of additional CD34^+^ and CD34^dim/−^neutrophil-restricted progenitors. (A) BM-LDCs were gated as SSC^low^CD45^dim^CD38^+^CD10^−^CD123^dim/−^ cells and then displayed based on their CD34 and CD45RA expression, in turn evidencing CD34^+^CD45RA^−^ (panel IV), CD34^dim/−^ CD45RA^+^ (panel VI), and CD34^dim/−^CD45RA^−^ (panel VII) cell populations [in addition to CD34^+^CD45RA^+^/cGMPs (panel V)]. Analysis of their cell composition revealed, other than those identified within cGMPs (green gate in panel V), the presence of CD34^+^CD45RA^+^CD64^dim^CD115^−^ populations, herein depicted as orange, magenta and light blue dots, and collectively renamed as NMs. All cell populations are overlapped in panel VIII. One representative experiment of at least 15 is shown. (B) Morphology of purified NM populations and PMs. Red arrows point to visible nucleoli, while blue arrows point to granules. (C) Bar graphs depicting the fold expansion of purified NM populations after a 7 day-culture with SFGc (mean±SEM of live, CD45^+^ output cells, n=6; *p < 0.05; **p < 0.01; ***p < 0.001). (D) Representative plots displaying the phenotype and morphology of CD66b^+^ neutrophils derived from NMs cultured with SFGc for 7 days. Bar graphs report the percentages relative to live, CD45^+^ cells (mean±SEM, n=6, *p < 0.05) of the various neutrophil lineage progenitors generated by NMs.

Collectively, these experiments formally prove that CD34^+^CD45RA^+^CD64^dim^CD115^−^ cells represent neutrophil-restricted progenitors. Data also demonstrate that the CD34^+^CD45RA^+^CD64^+^CD115^+^ cMoP-like and the CD34^+^CD45RA^+^CD64^−^CD115^+^ MDP-like cells substantially resemble, respectively, the conventional cMoPs ^17^ and MDPs ^16^, and thus they will be referred to as such.

### Identification of additional CD34^+^ and CD34^dim/−^neutrophil-restricted progenitors

Since neutrophil-committed progenitors have been suggested to stand within the CD45RA^−^ compartment ^3, 20^, we then wondered whether other, not yet described, neutrophil-restricted progenitors could be identified within the SSC^low^CD45^dim^ region (panel **I** of **Figure 2A**). Therefore, we searched for cells expressing the CD64^dim^CD115^−^ phenotype in all the fractions delimited by the CD34/CD45RA marker combination (panel **III** of **Figure 2A**), and ultimately found three more cell populations (orange, magenta and light blue gates in, respectively, panel **IV, VI** and **VII** of **Figure 2A**). Moreover, we confirmed the presence not only of the previously described CD34^+^CD45RA^+^CD64^dim^CD115^−^ neutrophil-restricted progenitors (green gate), cMoPs and MDPs within cGMPs (panel **V** of **Figure 2A**), but also of CD64^++^CD115^+^ pre-monocytes ^17^ in CD34^dim/−^ CD45RA^+^ cells (panel **VI** of **Figure 2A**). Notably, by applying an identical flow cytometry gating strategy to either cord blood samples (CB) or spleen biopsies, we were able to identify SSC^low^CD45^dim^CD123^dim/−^CD10^−^CD34^+^ and CD34^dim/−^ cells, as well as to successfully uncover, even in these specimens, CD64^dim^CD115^−^neutrophil-restricted progenitors within the CD45RA^+^ and CD45RA^−^ cells (**data not shown**). Hence, we renamed these CD34^+^ and CD34^dim/−^ neutrophil-restricted progenitors as neutrophil myeloblasts (NMs), and subdivided them, on the basis of their CD34 and CD45RA expression, as NM1s (orange gate in panel **IV**), NM2s (green gate in panel **V**), NM3s (magenta gate in panel **VI**) and NM4s (light blue gate in panel **VII**) for, respectively, CD34^+^CD45RA^−^, CD34^+^CD45RA^+^, CD34^dim/−^CD45RA^+^ and CD34^dim/−^CD45RA^−^ cells.

Then, we sorted CD34^+^ and CD34^dim/−^NMs to analyze their morphology and developmental potential. Morphologically (**Figure 2B**), the four NMs were found to display a high nuclear/cytoplasmic ratio compatible with a progenitor identity. Moreover, granules (**Figure 2B**) were evident only in NM3s and NM4s, but at lower levels than in PMs. In addition, SFGc-treated NM1s and NM2s were found to display a comparable (**Figure 2C**), but higher proliferative potential than that exhibited by SFGc-treated NM3s and NM4s (**Figure 2C**). Notably, the four NMs were found to almost exclusively produce neutrophils at different stages of maturation, as determined by their phenotype and morphology (**Figure 2D**). By the latter criteria, in fact, we found no eosinophils and/or basophils among the SFGc-generated cells (**data not shown**), and even if the four NMs were incubated in SFG plus IL-3 and/or IL-5 ^18^. Moreover, all NMs generated no DCs when cultured in SFGc (**data not shown**), while exclusively NM2s and NM3s generated a few CD14^+^ monocytes (**data not shown**), which we subsequently found out to actually derive from contaminating unavoidably sorted cMoP and pre-monocyte populations (see paragraph on single cell RNA data). Importantly, neutrophils generated by NMs displayed respiratory burst ability (**Figure S3A**), as well as phagocytosis capacity (**Figure S3B**) comparable to those displayed by peripheral mature neutrophils. Finally, we found that, unlike NM1s and NM2s, NM3s and NM4s express CD15, although at lower levels than PMs and mature SNs (**Figure S3C**). The latter data exclude that NMs could relate to the SSC^int^Lin^−^CD34^−^CD15^int^CD11b^−^CD16^−^ “early” PMs (EPMs) ^31^. In our hands, in fact, EPMs were found to partially overlap not only with both NM3s and NM4s, but also with pre-monocytes (**data not shown**), which express CD15 at similar levels of those by cMoPs and mature monocytes (**Figure S3D**). It is hence evident that that the sole CD15 does not function as a marker specifically identifying neutrophil progenitors.

In sum, our experiments have identified CD34^+^ and CD34^dim/−^NMs standing prior PMs along the neutrophil maturation cascade, and specifically characterized by the CD64^dim^CD115^−^ phenotype, but differentially expressing CD45RA. The features of CD34^+^ and CD34^dim/−^NMs are summarized in **Table S1**.

### The relative content of NMs, MDPs, cMoPs and CDPs is altered in chronic-phase chronic myeloid leukemia (CP-CML) patients

CML is a myeloproliferative disorder evolving in three clinical stages, known as CP-CML, accelerated phase CML (AP-CML) and blast phase CML (BP-CML) ^32^. CP-CML stands out for a dramatically increased peripheral white blood count (WBC) (i.e., higher than 10^10^/L), which mainly reflects a remarkable rise of both the absolute number and the relative percentage of neutrophils. Inexplicably, a dramatic decrease of the cGMP percentage has been reported to occur in CP-CML patients ^33-35^, which is in contrast with the classical model of myeloid development centered on cGMPs as the main source of neutrophils.

To try clarifying such an issue, we set up a 14-color antibody panel allowing a more clear identification not only of the SSC^low^Lin^−^CD45^dim^ region, but also of the various neutrophil progenitors, starting from their earliest CD34^+^ ones, and up to mature, segmented neutrophils (**Figure S4A and S4B**). This approach turned out to be fundamental to analyze CP-CML pathological samples. By this analysis, we confirmed that cGMPs from CP-CML patients decrease (**Figure 3A**) and result almost negative for CD115 expression (**Figure 3B**), therefore reflecting an altered ratio between neutrophil and mono-DC progenitors (**Figure 3C**). Precisely, we found the percentage of MDPs, cMoPs and CDPs as significantly lower in CP-CML patients than in HDs (**Figure 3C**), in the presence of a significant expansion of NM1s and of an unaltered NM2 frequency (**Figure 3C**). Therefore, while confirming the presence of NMs in CML patients, these data imply that the down-regulation of cGMP percentage in CP-CML patients depends on a marked reduction of monocyte/DC progenitors but not of NM2s. Data also support the notion that the selective expansion of neutrophils in CP-CML patients derives from NM1s.

**Figure 3.**
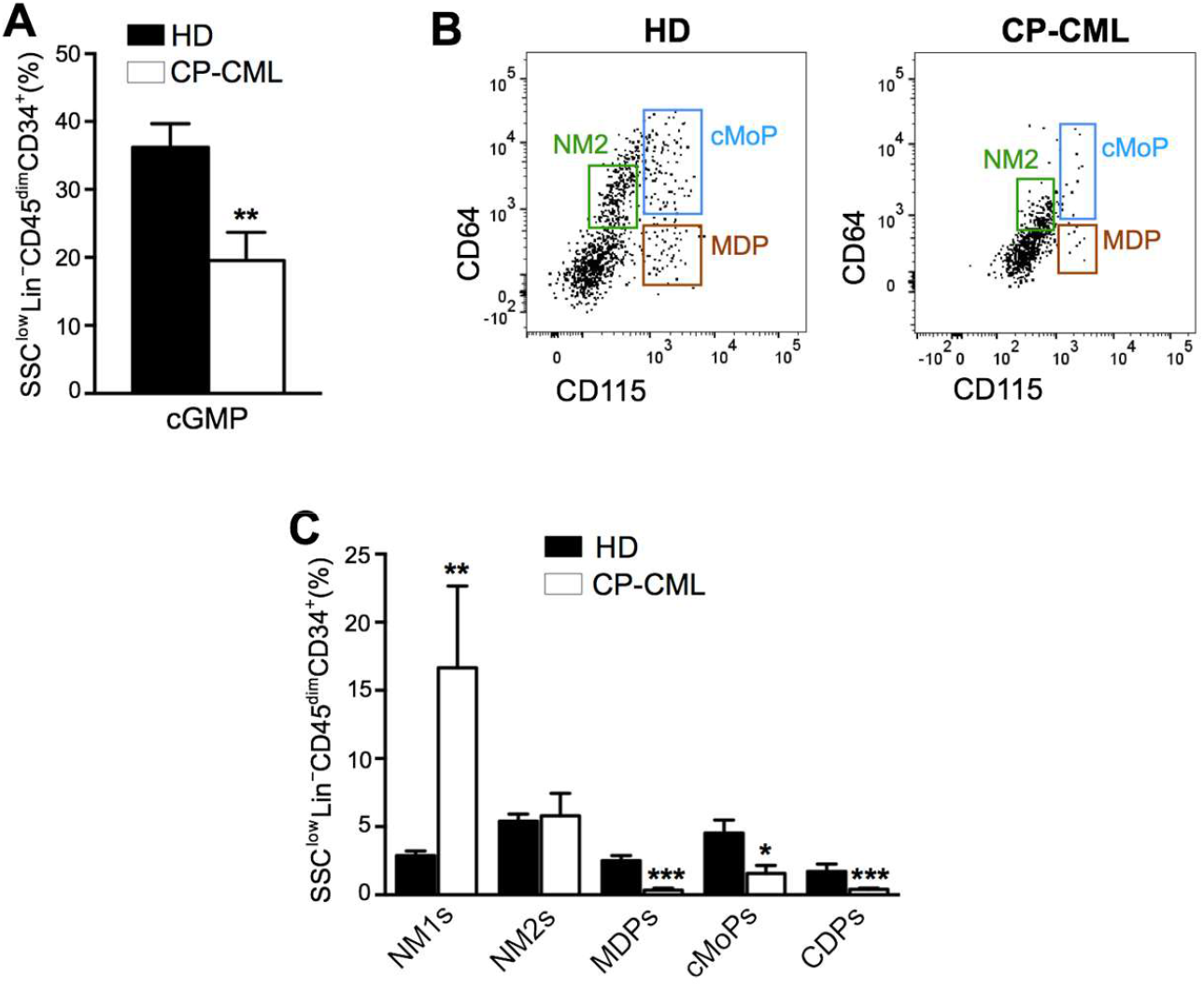
The relative content of NMs, MDPs, cMoPs and CDPs is altered in chronic-phase chronic myeloid leukemia (CP-CML) patients. BM-LDCs from HDs (black bars) and CP-CML patients (white bars) were subjected to the 14-color gating strategy illustrated in Figure S4. (A) Bar graphs showing the percentage of cGMPs relative to SSC^low^Lin^−^CD45^dim^CD34^+^ cells (mean±SEM, n=7, **p < 0.01). (B) Representative flow cytometry plots showing the distribution of NM2s, cMoPs and MDPs. (C) Bar graphs showing the percentage of NM1s and NM2s, as well of MDPs, cMoPs and CDPs, relative to SSC^low^Lin^−^CD45^dim^CD34^+^ cells (mean±SEM, n=7, *p < 0.05; **p < 0.01; ***p < 0.001).

### RNA-seq experiments confirm that NMs precede PMs along the neutrophil maturation cascade

NMs, as well as BM HSCs, PMs, MYs, MMs, BCs, SNs and mature neutrophils (PMNs) were then profiled by RNA-seq. Both PCA (shown in **Figure 4A**) and hierarchical clustering analysis performed by optimal leaf ordering (OLO) ^36^ (**Figure 4B**) confirmed that NMs not only cluster by themselves, but are also placed along the maturation trajectory which from HSCs, *via* PMs, MYs, MMs, BCs and SNs, ends into PMNs. Interestingly, analysis of *CD34* and *PTPRC* (encoding for CD45) transcript expression (**Figure 4C** and **4D**) corroborated the flow cytometry data. Furthermore, by performing K-means clustering of the differentially expressed genes (DEGs), ten main gene groups (g1-g10) were identified among all samples (**Figure 4E** and **Table S2**). Genes encoding markers of immature cells (such as *CD34, HOXA9, MYC, FLT3, SOX4* and *KIT*), expressed only by HSCs and NMs, were found present in g1 or g2 (**Figure 4E**). Interestingly, GO analysis of the g1 and g2 DEGs revealed the “MHC class II protein complex” term among the most significant over-represented (**Figure S5A**), in line with a previously described MHC class II expression in very immature neutrophil progenitors ^37^. Similarly, DEGs mainly involved in ribosome assembly, mitochondria formation and cell cycle regulation, characterizing g3, g4 and g5, respectively, were found expressed in NMs, PMs, MYs and MMs, but not in non-proliferating BCs and SNs (**Figure 4E**,**F** and **Figure S5A**).

**Figure 4.**
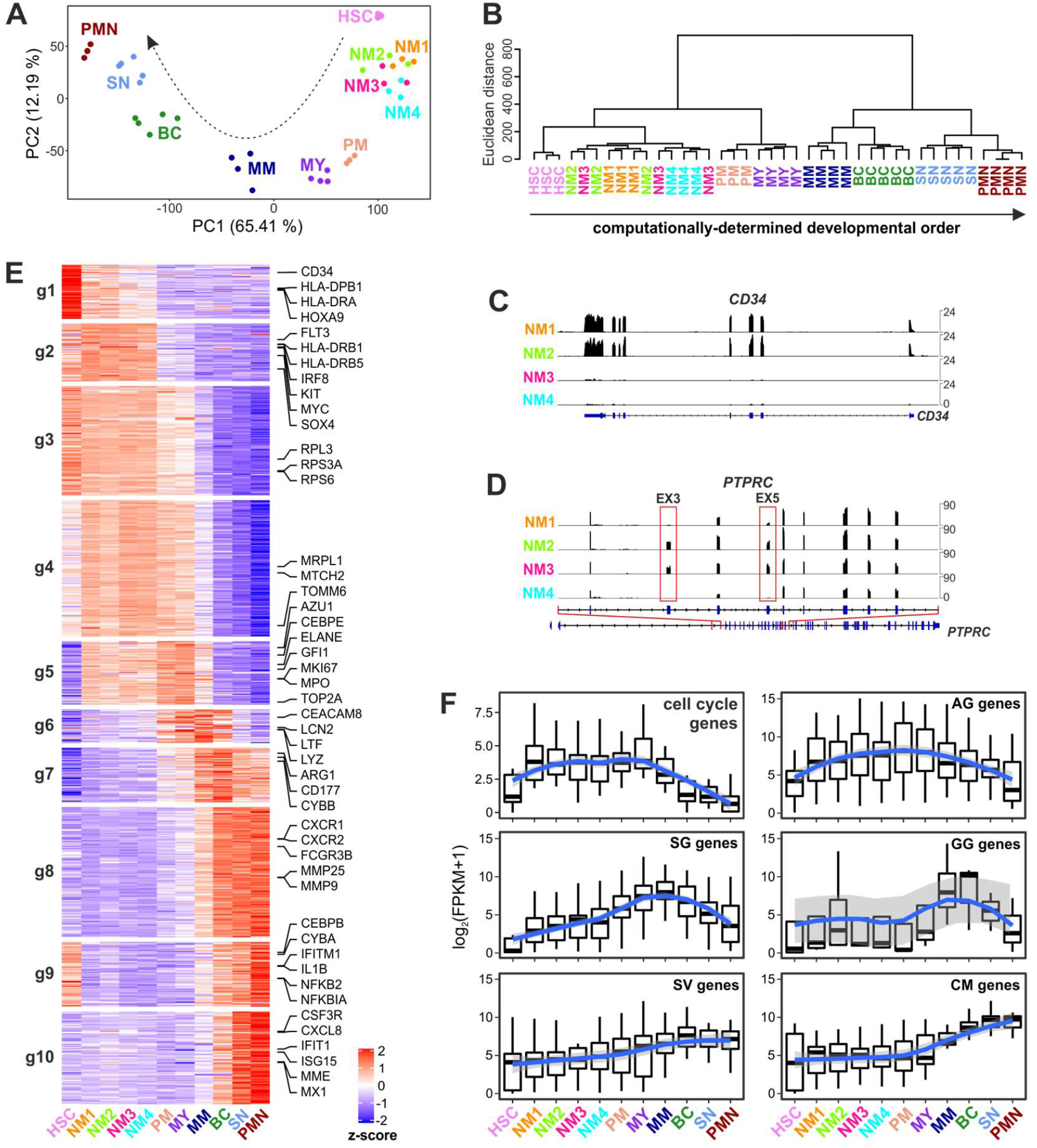
RNA-seq experiments confirm that NMs represent very early precursors of neutrophils. (A) PCA scatter plot based on the DEGs identified from bulk RNA-seqs of NM1s, NM2s, NM3s and NM4s, as well as HSCs, PMs, MYs, MMs, BCs, SNs and mature neutrophils (PMN) (n= 3-5). (B) Developmental path of the various neutrophil lineage progenitors computationally determined from bulk RNA-seq datasets using the optimal leaf ordering (OLO) algorithm. (C, D) Scheme represent genome-browser views of RNA-seq signals at the *CD34* (C), and *PTPRC/CD45* (D), loci of NMs. Exons 3 and 5 of *PTPRC*, whose transcription is essential for the expression of the CD45RA variants, are highlighted by red boxes in (D). (E) Heatmap displaying the expression patterns of the gene groups (g1–g10) resulting from the K-means analysis of DEGs identified among the various neutrophil-lineage cells. The average gene expression levels of biological replicates were calculated, and data were represented as z score. (F) Box plots showing the distribution of the expression levels [as log2(FPKM+1)] of cell cycle, AG, SG, GG, SV and GM mRNAs across neutrophil differentiation. Upper and lower boxplot margins indicate first and third quartiles of genes expression levels. LOESS fitting of the data, with relative confidence interval, is represented by a blue line with a shadow area.

Importantly, g5 was also found to contain DEGs encoding the first granule mRNAs transcribed during granulopoiesis, namely those for the azurophilic granule (AG) proteins (such as MPO, AZU1, PRTN3 or ELANE), as well as transcription factors (TFs) known to be typically expressed in immature neutrophils, such as GFI1 and CEBPE ^38^ (**Figure 4E**). Focused/specific analysis on AG genes expression revealed that these genes start to be expressed in NM1s and NM2s, increase in NM3s and NM4s, are maximally transcribed in PMs and MYs and then gradually disappear (**Figure 4F**). By contrast, g6, g7 and g8 were found to include DEGs absent in NMs, being enriched for specific granule (SG) (LYZ, LTF, LCN2 and CEACAM8/CD66b), gelatinase granule (GG) (MMP9, ARG1 and CD177), secretory vesicles (SV) (e.g., FCGR3B/CD16B and ANXA1) and cell membrane proteins (CM) (e.g., CXCR1, CXCR2 and ICAM3) mRNAs (**Figure 4E**,**F**) and thus associated to “neutrophil degranulation” and “neutrophil activation” GO terms (**Figure S5A**). Finally, g9 and g10 were found enriched in IFN-stimulated genes (ISGs), as well as other genes associated to the “NF-kB signaling” and “cytokine production” GO terms, and all of them present in BCs, SNs and PMNs (**Figure 4E** and **Figure S5A**). Consistent with the literature ^39^, mRNAs encoding proteins involved in the production of reactive oxygen species (ROS) (**Figure S5B**), phagocytosis (**Figure S5C**) and chemotaxis (**Figure S5D**) were found transcribed starting from the MM/BC stages, and highly expressed in mature neutrophils.

To get more insights into the specific transcriptomic differences among NMs only, we performed DEG analysis by using the likelihood ratio test (LRT)^40^ and identified 1114 DEGs among them. As shown in **Figure S5E**, the four NMs were found to distinctly segregate from each other by PCA, indicating remarkable differences among their gene expression profiles. However, while PC1 differences were mostly determined by cell cycle and AG genes, which were more expressed in, respectively, NM1s/NM2s and NM3s/NM4s (**Figure S5E**), genes mostly contributing to the PC2 variations, namely *PRG2, CLC, EPX* and *IL5RA* mRNAs found in NM1s, or *IRF8, ANXA2, SAMHD1, SLAMF7, LYZ* and *F13A1* mRNAs found in NM2s and NM3s (**Figure S5E**), were unexpected since typically expressed by progenitors of, respectively, eosinophils or monocytes. All in all, RNA-seq data not only confirm that NMs are committed to neutrophils, but also that they are placed prior to PMs along the neutrophil maturation cascade.

### Identification of multiple cell clusters within NMs by scRNA-seq

To unequivocally dissect their transcriptional heterogeneity, we performed scRNA-seq experiments of sorted NM1s, NM2s, NM3s, NM4s, and (to elucidate their relationship with NMs) cMoPs. We sequenced 17902 cells and, by performing dimensionality reduction by Uniform Manifold Approximation and Projection (UMAP) ^41^, it immediately emerged that NMs clearly segregate from and cMoPs, while NM1s and NM3s mostly overlap with, respectively, NM2s and NM4s (**Figure 5A** and **Figure S6A**). Then, by performing unbiased, graph-based, clustering by Seurat ^42^, we could identify 9 discrete cell clusters (**Figure 5B**), of which three (c7, c8 and c9) were excluded from all subsequent analyses because consisting of: i) few eosinophil progenitors (i.e., c7, identified by *EPX* mRNA expression); ii) few megakaryocyte/erythroid progenitors (i.e., c8, identified by *PF4* and *HBB* mRNA expression); iii) few cells with an unclear ontogeny and high expression of apoptosis-related genes (i.e., c9) (**Figure 5B**). Of the six remaining cell clusters, by far the most rich of cells, four were unequivocally attributable to the neutrophil lineage (i.e., c1, c2, c3 and c4), and two to the monocyte lineage (i.e., c5 and c6) (**Figure 5B**). Such segregation was also confirmed by hierarchical clustering analysis of their DEGs (**Figure S6B**), since genes typical of the neutrophil (such as *CEBPE* or *GFI1*) and monocyte (such as *IRF8* or *CSF1R*) lineages were exclusively expressed in, respectively, c1-c4 and c5-c6 cells (**Figure 5C**). Moreover, distribution analysis of scRNA-seq cell clusters in NMs and cMoPs evidenced that c1-c4 cells were completely absent in cMoPs (thus proving that CD115 expression efficiently discriminates NMs). By contrast, a quote of c5 cells were found to contaminate both NM2s and NM3s, but not NM1s and NM4s (**Figure 5D**), therefore explaining the PCA results on bulk RNA-seq shown in **Figure S5G**.

**Figure 5.**
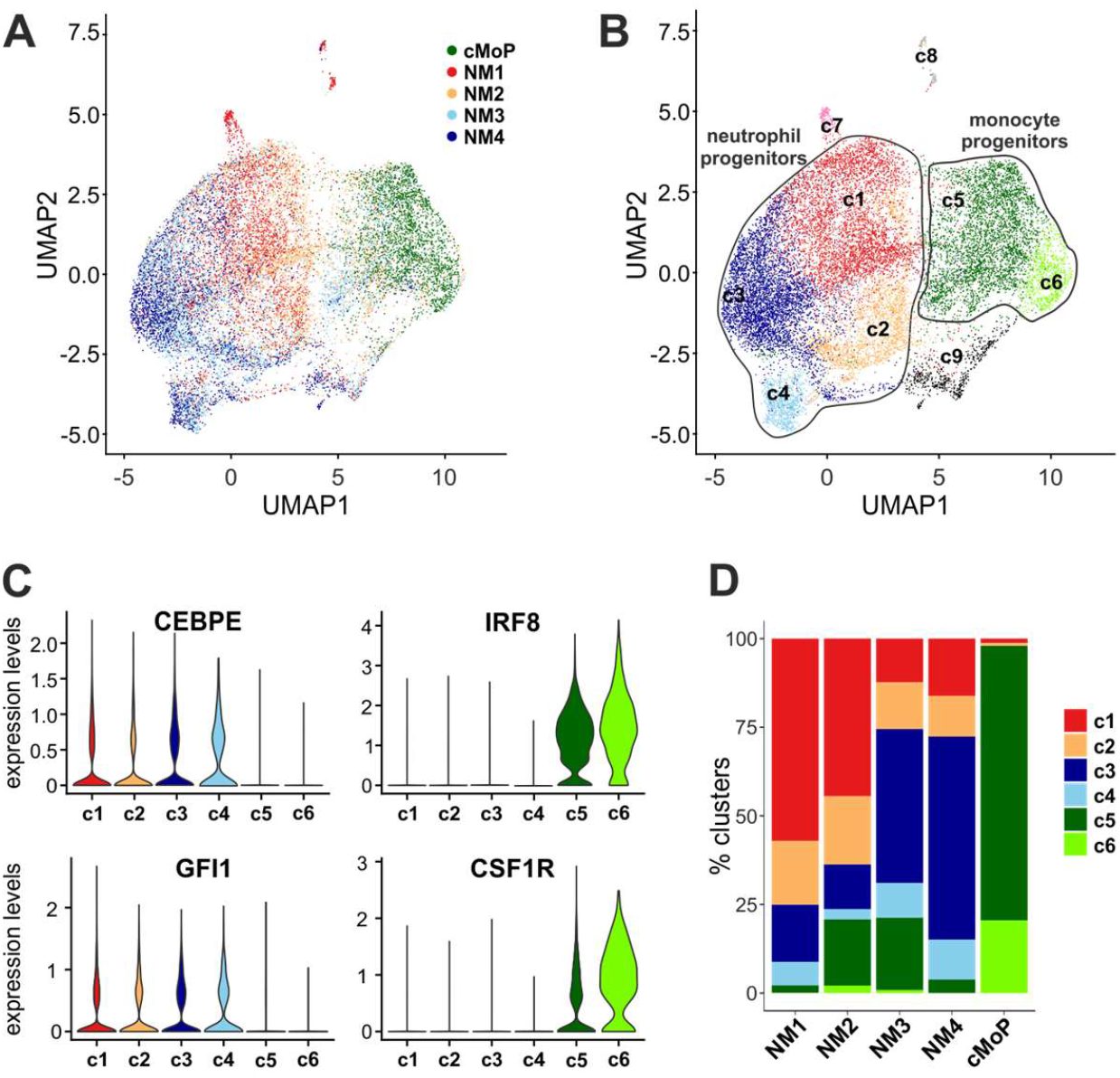
scRNA-seq experiments of NMs and cMoPs reveal that they consist of multiple cell clusters. (A) UMAP visualization of scRNA-seq profiles of NM1s, NM2s, NM3s, NM4s and, for comparison, cMoPs, isolated from 3 different BMs. Cells are colored according to the cell type identification shown in Figure 2A. (B) UMAP shown in (A) colored according to the inferred cell cluster identity (c1-c7), as identified by Louvain clustering. (C) Violin plots showing the mRNA expression levels of *CEBPE* and *GFI1* (neutrophil lineage-specific genes), and *IRF8* and *CSF1R* (monocyte lineage-specific genes), across c1-c6 cells. (D) Bar graph showing the relative abundance of c1-c6 cells in NM1s, NM2s, NM3s, NM4s and cMoPs.

By selectively focusing on c1-c4 cells, it resulted evident that their distribution in NM1s and NM4s is specular to those of, respectively, NM2s and NM3s (**Figure 6A**). Moreover, while NM1s and NM2s were found enriched of c1 cells mainly, NM3s and NM4s were found to preferentially accumulate c3 and c4 cells (**Figure 6A**). By contrast, c2 cells were found to distribute among the four NMs at substantially similar levels (**Figure 6A**). Then, calculation of the pseudotime value for every cell by Destiny ^43^(**Figure 6B**), to define their maturation state, uncovered that c1 contains the more immature, while c3 and c4 contain the relatively more mature, cells (**Figure 6C)**. Accordingly, *CD34* was found preferentially expressed in c1 and c2 cells (**Figure 6D**), consistent with their relatively more abundant presence in NM1s and NM2s (**Figure 6A**). c2 cells, instead, were found to display variable pseudotime values (**Figure 6C**), in line with their presence in all NMs (**Figure 6A)**. Assessment, by Seurat, of the DEGs characterizing c1-c4 cells resulted in the identification of 136 genes (**Figure 6E** and **Table S3**). In such regard, c1 cells were found to express high levels of typical genes of immature proliferating cells (such as *CD34, TOP2A, TUBB* and *HIST1H4C*) (**Figure 6E** and **Figure S6C**,**D**), consistent with their lowest pseudotime values (**Figure 6C**). c1 cells were found to express also genes associated to the “MHC class II protein complex” GO term (**Figure S6E)**, consistent with the g2 genes from bulk RNA-seq data (**Figure S5A**). By contrast, c3 and c4 cells were found to express high levels of AG genes (**Figure 6E** and **Figure S6C**,**D**), as well as to display GO terms mostly enriched for “neutrophil activation” and “neutrophil degranulation” (**Figure S6E**), in accordance with their elevated pseudotime values (**Figure 6C**), and their prevalent correspondence with NM3s and NM4s (**Figure 6A**). Notably, c4 cells were also found to specifically express high mRNA levels of *BEX1* (**Figure 6E** and **Figure S6C**), while unexpectedly, but in in line with “defense response to virus” as the GO term most enriched for them (**Figure S6E**), c2 cells were found to express elevated levels of interferon-stimulated genes (ISGs), such as *ISG15, IFI6, IFIT3* and many more (**Figure 6E** and **Figure S6C**). No SG or GG genes were present in scRNA-seq datasets (**Figure S6D**), confirming RNA-seq data (**Figure 4D**). Finally, by assessing their potential developmental trajectories [by using Destiny algorithm ^43^], we found that c1-c4 cells distribute along three branches, one of them including the majority of c1 cells (**Figure 6F**,**G** and **Figure S6F**) other than a minor quote of the c2, c3 and c4 cells. The c1 branch was then found to continue into two different trajectories (**Figure 6F)**: one, defined as “conventional trajectory”, since mainly characterized by cells expressing elevated AG mRNA levels (i.e., c3 and c4 cells), but also including the *BEX1* mRNA-positive cells (c4 cells); the other one, unexpected, and defined as “ISG trajectory” (**Figure 6F**) since characterized by cells mainly expressing high ISG mRNA levels (i.e., c2 cells) (**Figure 6H**). In sum, scRNA-seq clustering analysis of NMs identified four clusters of neutrophil progenitors at different stages of maturation and distributed along two maturation routes. Data also uncovered that, even though phenotypically differing among themselves exclusively at the CD45RA level, NM1s and NM2 on the one hand, and NM3s and NM4s on the other hand, are identical in terms of cluster distribution.

**Figure 6.**
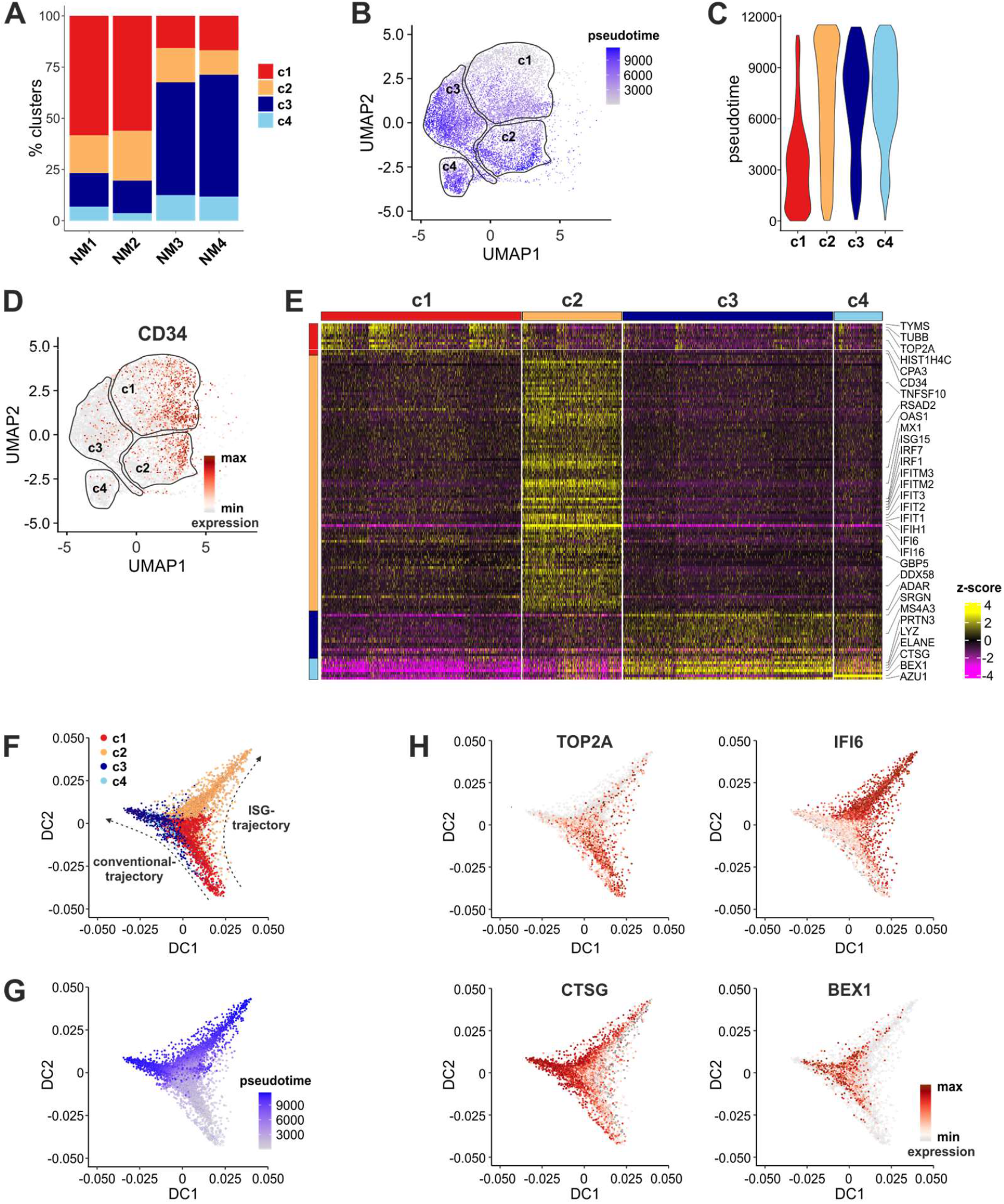
Characterization of the scRNA-seq cell clusters composing the NMs. (A) Bar graph showing the relative abundance of c1-c4 cells in NM1s, NM2s, NM3s and NM4s, after exclusion of cells from monocyte progenitors (c5 and c6). (B,C) UMAP (B) and violin (C) plots restricted to c1-c4 cells, showing the pseudotime value for every cell, as calculated by Density algorithm. (D) Expression patterns of *CD34* mRNA projected on a UMAP plot restricted to c1-c4 cells. (E) Heatmap showing scaled expression [log TPM (transcripts per million) values] of discriminative gene sets (Bonferroni-corrected P values < 0.05; Student’s t-test) by c1-c4 cells, as defined in Figure 5B. Color scheme is based on z-score distribution from –4 (purple) to 4 (yellow). (F,G) Trajectory plots showing the distribution of c1-c4 cells (F), and their pseudotime values (G). In F, the arrows indicate both the “conventional”, and the “ISG”, developmental trajectories, along the neutrophil lineage. (H) mRNA expression patterns of genes characteristic of c1 (i.e., *TOP2A*), c2 (i.e., *IFI6*), c3 (i.e., *CTSG*) and c4 (i.e., *BEX1*) cells, projected on the trajectory plots from (F).

## DISCUSSION

In this study, by using the CD45/CD38/CD34/CD10/CD123/CD45RA/CD64/CD115 flow cytometry antibody panel to examine the SSC^low^CD45^dim^ region of human BM, we initially identified CD34^+^CD45RA^+^CD64^dim^CD115^−^ cell progenitors, within cGMPs, exclusively committed to the neutrophil lineage. These neutrophil progenitors may therefore be the long searched-for “missing piece in the puzzle” ^44^ that composes the cGMP region together with the uni-potent cMoPs and CDPs, and the heterogeneous MDPs and GMDPs. By the same approach, not only we confirmed cMoPs as CD64^+^CD115^+^ ^17, 21^, but also pointed both MDPs and GMDPs as CD64^−16^. Since our data are in line with results from other groups ^4, 18-20^, we propose to rename GMDPs as NMDPs (i.e., neutrophil-monocyte-dendritic cell progenitors).

Subsequent flow cytometry and *in vitro* differentiation experiments uncovered, other than those initially found within cGMPs, three more CD34^+^ and CD34^dim/−^CD64^dim^CD115^−^ neutrophil progenitors, which we ultimately renamed as neutrophil myeloblasts (NMs), and based on their differential CD45RA expression, subdivided into NM1s for CD34^+^CD45RA^−^ NMs, NM2s for CD34^+^CD45RA^+^ NMs, NM3s for CD34^dim/−^CD45RA^+^ NMs and NM4s for CD34^dim/−^CD45RA^−^ NMs. It is correct to point out that, since the CD34 and CD45RA fluorescence distributions in SSC^low^CD45^dim^ cells change gradually during their maturation, the separation of CD34^+^ and CD34^dim/−^ NMs into four phenotypical populations has been done to intentionally enrich for cells at similar maturation stages. However, since hematopoietic progenitors flow in continuum during their differentiation process, it is implicit that also NMs mature *via* gradual transitions at both phenotypical and transcriptional levels. The biological significance of the presence of both CD45RA^+^ and CD45RA^−^ NMs remains to be clarified. One hypothesis could be that CD45RA serves to localize CD34^+^ and CD34^dim/−^NMs in specific BM niches. Another one could be that CD45RA has a functional, yet unknown, role under discrete pathological conditions, as suggested by the selective expansion of CD45RA^+^ immature cells in the majority of acute myeloid leukemia patients ^45^.

That CD34^+^ and CD34^dim/−^NMs represent very early progenitors has been unequivocally confirmed by RNA-seq experiments, which indeed proved that they do not express SG genes. Moreover, clustering analysis of DEGs among CD34^+^ and CD34^dim/−^NMs evidenced a remarkable enrichment of genes related to cell proliferation in NM1s/NM2s, and of AG-encoding genes in NM3s/NM4s. These findings exclude that our CD34^+^ and CD34^dim/−^NMs correspond to the previously described human preNeus ^22^, which instead express high levels of SG genes. NM2s, instead, might eventually correspond to the recently described murine proNeu1s, defined as early committed neutrophil progenitors preceding preNeus found within GMPs and not expressing SG genes ^9, 24^. However, the human counterpart of proNeus, identified by “Infinity Flow” in cord blood and fetal BM ^24^, was found to be CD66b^+^/CD15^hi^, unlike our CD34^+^ and CD34^dim/−^NMs. Moreover, our NMs, other than being CD66b^−^, are included within the SSC^low^CD45^dim^ progenitor region of BM, which proves that they represent neutrophil progenitors more immature than the recently described CD66b^+^ hNePs ^23^, eNePs ^25^, or proNeus described in severe COVID-19 patients ^46^.

scRNA-seq experiments further uncovered that the CD34^+^ and CD34^dim/−^NMs actually consist of four cell clusters (i.e., c1-c4), displaying distinct features (see results), and distributed at substantially identical ratios in NM1s and NM2s, as well as in NM3s and NM4s. Intriguingly, c4 cells were found to specifically express elevated mRNA levels of BEX1, a protein reported to function as a tumor suppressor in FLT3-ITD positive AML ^47^, as well as to impact on the response to Imatinib in CML patients ^48^. Moreover, by inference methods, we revealed that the four cell clusters distribute along two developmental trajectories originating from c1 cells, namely one, involving c3/c4 cells, that we indicated as “conventional” for the expression of genes characteristic of the neutrophil lineage; the other one, involving c2 cells, that we defined as “ISG” trajectory for their transcriptomic features. The existence of neutrophil progenitors enriched in ISG genes is in line with the recent identification of mature neutrophil subsets expressing ISGs under either healthy ^49, 50^ or pathological ^46, 51, 52^ conditions. Intriguingly, recent scRNA-seq data have shown that ISGs are enriched in BCR-ABL mutation-free HSCs of CP-CML patients, who poorly respond to Imatinib treatment ^53^.

Clusters of neutrophil progenitors, displaying levels of maturation apparently similar to our c1-c4 cells, have been also identified by scRNA-seq studies of HSPCs from human BM, even though no sorting strategy for their isolation was described. For instance, a cell cluster of myeloid progenitors expressing *ELANE* and *PRTN3* has been found within CMPs ^20^, but, as such, it might be also confused with cells belonging to the monocyte lineage. In another of these studies ^3^, four clusters of neutrophil progenitors (i.e., N0-N3) were found in CMPs (i.e., N0) and cGMPs (i.e., N1, N2 and N3). N1, N2 and N3 might in part correspond to our c1 cells, given their CD34 protein, as well as AG mRNA, expression. However, N3 could be rather ascribed to a monocyte progenitor for its expression of *LYZ, SAMHD1, CSF1R, ANXA2, KLF4* and *IRF8* genes ^3^, which we found to be peculiar to cMoPs (**our unpublished data**). Similarly, since shown to include only MPO among AG genes ^3^, which can be also shared by cMoPs ^9^, it remains uncertain whether N0 effectively corresponds to a neutrophil progenitor. None of these studies ^3, 20^ detected specific ISG- or BEX1-containing clusters, likely for the much lower number of neutrophil progenitors investigated than in our scRNA-seq set.

In summary, we identified CD34^+^ and CD34^dim/−^ neutrophil-committed progenitors within human BMs, all of them characterized by a SSC^low^Lin^−^CD45^dim^CD64^dim^CD115^−^ phenotype, that in our view correspond to neutrophil myeloblasts. As such, CD34^+^ and CD34^dim/−^NMs are easily sortable and manageable for further studies. We expect that future work could uncover specific membrane markers allowing the sorting of the cell clusters composing CD34^+^ and CD34^dim/−^NMs, to perform a better characterization and also to evaluate if and how they expand under pathological conditions.

## ACKNOWLEDGMENTS

This work was supported by grants to MAC (from AIRC IG-20339; MIUR-PRIN 20177J4E75_004; and Fondazione Cariverona) and to N.T (GR-2016-02361263). We thank the Centro Piattaforme Tecnologiche (CPT) of University of Verona for the access to the flow cytometry/cell analysis and genomic/transcriptomic platforms.

## AUTHOR CONTRIBUTIONS

F.C. and G.F. performed and analyzed flow cytometry, cell sorting and *in vitro* differentiation experiments; C.C. and A.M. were involved in cell sorting experiments; N.T., M.C. and S.G. prepared samples for RNA-seq and scRNA-seq; N.T. and F.B.A. analyzed RNA-seq and scRNA-seq data; F.B., M.B. and C.T. provided patient samples; F.C., G.F., N.T., P.S. and M.A.C. designed and wrote the manuscript; M.A.C supervised the project.

## METHODS

### Isolation of cells from human bone marrow (BM) and peripheral blood samples

Upon local ethical committee approval, BM samples were obtained from healthy donors (HDs) (n=29, Table S4) selected for BM donation purposes according to the Italian Bone Marrow Donor Registry (IBMDR) criteria. 1.5/2 ml fresh BM samples were collected under aseptic conditions in heparinized sterile syringe, processed using endotoxin-free polypropylene tubes (Greiner bio-one), and subjected to density gradient centrifugation onto Ficoll-Paque (GE Healthcare). BM low-density cells (BM-LDCs) were then collected, and either immediately processed for flow cytometry analysis, or suspended in αMEM growth medium (Corning) supplemented with 10 % low-endotoxin FBS (< 0.5 EU/ml), in the presence of penicillin/streptomycin (from now on termed “culture medium”)^29^, to be finally distributed in tissue-culture plates (Corning) and preincubated for 20 h at 37° in CO2 ^21^. Occasionally, fresh aliquots of BM-LDCs were frozen in 90 % FBS (Sigma-Aldrich) plus 10 % DMSO (Sigma-Aldrich) and stored in liquid nitrogen, since results obtained by using thawed or unfrozen BM-LDC samples were found fully comparable. Neutrophils from HDs were purified by the EasySep Human Neutrophil Enrichment Kit (Stem Cell Technologies; > 99.7% purity) after centrifuging buffy coats onto Ficoll-Paque gradient density, as described ^54^. BM samples from CP-CML patients (n=7, whose features are listed in Table S5) were obtained from subjects at first CML diagnosis without treatment (including hydroxyurea). CML diagnosis was made according to current standards ^55, 56^.

### Study approval

Human samples were obtained following informed written consent by HDs and CML patients. The study has been cleared by the Ethic Committee of the Azienda Ospedaliera Universitaria Integrata di Verona (Italy) (CMRI/55742).

### Flow cytometry and fluorescence-activated cell sorting experiments

For flow cytometry experiments, BM-LDCs, sorted progenitor subpopulations or culture-derived cells were counted, resuspended in 50 µl PBS buffer (Corning) plus 2 % FBS and 2 mM EDTA (Sigma-Aldrich) (from now on termed “staining buffer”), and subsequently incubated for 10 min in the presence of 5 % human serum (Sigma-Aldrich). Cells were then stained for 30 min on ice by fluorochrome-conjugated monoclonal antibodies (mAbs) for either the 8-color MACSQuant Analyzer (Miltenyi Biotec), or the 14-color LSRFortessa™ X-20 flow-cytometers (BD), as listed in Table S6. Viability of both fresh and 20 h-preincubated BM-LDC samples was assessed by either PI (Sigma-Aldrich), or alternatively Sytox Blue (Invitrogen) exclusion, and reproducibly found > 95 % (data not shown). The mean fluorescence intensity (MFI) relative to selected markers was obtained by subtracting either the MFI of the correspondent isotype control mAbs, or the cell autofluorescence (FMO), to their fluorescence value. FlowJo software version 10.7.1 was used for data analysis. For fluorescence-activated cell sorting of the myeloid progenitors, 50-100*106 BM-LDCs were resuspended at 100*106/ml and labeled for 45 min at 4° (in the dark) with the mAbs reported in Table S6. Anti-CD14 and anti-IL5Rα mAbs were always included in the sorting panel to exclude, respectively, mature monocytes and eosinophil-committed progenitors. Conversely, we omitted the use of antiCD38 and antiCD117 antibodies since NMs were found as CD38^+^ and CD117^+^. Cells were then washed and resuspended at 30*10^6^/ml in staining buffer, to be ultimately filtered through a 0.35 µM nylon mesh. Cells were finally sorted by using a FACSAria Fusion™ (BD) cell sorter equipped with 85-μm nozzle, immediately centrifuged, resuspended in αMEM medium, counted and used for experiments. Alternatively, sorted cells were lysed in RLT buffer (Qiagen) for RNA extraction ^57^. Sorted cell populations displayed a > 95 % purity, as verified by flow cytometry analysis.

### *In vitro* differentiation assay

To evaluate the differentiation potential of sorted BM progenitors, the latter cells were cultured on the MS-5 stromal cell line (from Leibniz-Institut DSMZ Braunschweig), according to protocols previously described ^29^. Briefly, the day before the coculture experiments, MS-5 cells cultured in αMEM medium at 95 % confluence were incubated with 10 µg/ml mitomycin C (Sigma-Aldrich) for 3 h at 37°. Then, after treatment with 0.05 % trypsine/EDTA (Corning), MS-5 cells were harvested and resuspended in αMEM medium at 0.25*10^6^/ml, to be finally seeded in round bottom 96-well tissue-culture plates for 24 h. 1-3*10^3^ of the various sorted BM progenitors were thus resuspended in 100 μl αMEM medium, seeded on top of MS-5 cells, and incubated with 20 UI/ml Flt3L (Miltenyi Biotec), 10 UI/ml SCF (Miltenyi Biotec), and either 100 UI/ml GM-CSF (Peprotech)^29^ or, as in previous studies ^58, 59^, 6500 UI/ml G-CSF (Myelostim, Italfarmaco Spa), to compose, respectively, the SFG ^29^ or SFGc cocktail. Cells were then harvested from the MS-5 cells starting at day 1, and up to day 14, of the culture, either for flow cytometry staining, or for functional studies. Expansion of purified progenitor populations is reported as mean fold-change of live, CD45^+^ output cells from the number of cells plated as input at time 0.

### Morphological analysis of neutrophil progenitors and culture-derived cell populations

Cytospins prepared from either sorted neutrophil progenitors or culture-derived cell populations were stained by the May-Grunwald Giemsa procedure. Slides were analyzed by a Leica DM 6000 B microscope, equipped with a Leica DFC 300FX Digital Color Camera (Leica Microsystem Wetzlar, Germany).

### O_2_^−^ production and phagocytosis assay

Either progenitor-derived neutrophils harvested from NMs cultured for 7 days, or freshly isolated HD neutrophils, were washed and resuspended at 0.25*10^6^/ml in HBSS supplemented with 10 % FBS, containing 1 mM CaCl_2_ and 5 mM glucose. O_2_^−^production in response to 20 ng/ml PMA (Sigma-Aldrich) was assessed by the Cytochrome C reduction assay, as previously described ^60^. To assess phagocytosis capacity, 0.025*10^6^ cells/100 µl were incubated in Eppendorf tubes at 37°, in the absence/presence of 20 µg/ml unopsonized zymosan particles. Cells were intermittently resuspended and, after 30 min, treated with an excess of cold HBSS and centrifuged to stop phagocytosis. Cytospin preparations of these samples were stained by the May-Grunwald Giemsa procedure, and slides analyzed by a Leica DM 6000 B microscope, equipped with a Leica DFC 300FX Digital Color Camera (Leica Microsystem Wetzlar, Germany).

### RNA sequencing (RNA-seq)

Total RNA was extracted by the RNeasy Mini Kit (Qiagen, Venlo, Limburg, Netherlands) after cell lysis ^61^. To completely remove any possible contaminating DNA, an on-column DNase digestion with the RNase-free DNase set (Qiagen) was performed during total RNA isolation ^62^. Quality control of the total RNA was performed using Agilent TapeStation 2200 (Agilent Technologies). RNA integrity (RIN) was routinely found to be optimal (RIN ≥ 7.0). Libraries for transcriptome analysis were prepared using the Smart-seq2 protocol ^63^. Briefly, 2 ng total RNA were copied into a first strand cDNA by reverse transcription and template-switching, using oligo (dT) primers and a LNA-containing template-switching oligo (TSO). The resulting cDNA was pre-amplified, purified, and tagmented with Tn5 transposase (kindly gifted by Dr. Sebastiano Pasqualato, European Institute of Oncology, Milan, 20139, Italy). cDNA fragments generated after tagmentation were gap-repaired, enriched by PCR and purified to create the final cDNA library. Libraries were sequenced on the Illumina NextSeq 500 in single-read mode (1×75 cycles) at the Centro Piattaforme Tecnologiche (CPT) of the University of Verona.

### Single cell RNA sequencing

Sorted cells were labelled by using the BD Single-Cell Multiplexing Kit (BD Biosciences), strictly following the manufacturer’s protocol (BD Biosciences). Such a procedure allowed us to combine 5 samples (NM1s, NM2s, NM3s, NM4s and cMoPs) into a single pool. Each sample was washed twice in FACS buffer and resuspended in cold BD Sample Buffer (BD Biosciences). Only samples with high viability (> 85 %, as evaluated by the BD Rhapsody scanner) were used for sequencing. Samples from the same donor were pooled in equal amounts to achieve approximately 15000 cells in 620 µl, and loaded onto a BD Rhapsody cartridge for an incubation of 20 min at room T. Then, Cell Capture Beads (BD Biosciences) were added to the cartridge, incubated at room T for 3 min, and thereafter Cartridges washed. Cell were then lysed, and the released mRNA captured by Cell Capture Beads. mRNA were then retrieved, to be washed prior to performing reverse transcription and treatment with Exonuclease I. cDNA Libraries were prepared by using the BD Rhapsody Whole Transcriptome Analysis (WTA), Amplification kit and BD Single-Cell Multiplexing kit (BD Biosciences). Size-distribution (Quality) of final libraries was assessed by Agilent 2200 TapeStation with High Sensitivity D5000 ScreenTape and quantified using a Qubit Fluorometer using the Qubit dsDNA HS Kit (ThermoFisher, #Q32854). Sequencing was performed in paired-end mode (2×75 cycles) on a NextSeq 500 System (Illumina). This procedure was utilized for three different BMs, whose data were integrated as outlined below.

### RNA-seq computational analysis

Computational analysis of transcriptome datasets generated by Smart-seq2 has been performed by using the bioinformatic pipeline, as previously described ^64^. Briefly, after quality filtering, according to the Illumina pipeline, removal of contaminant adapters and base quality trimming were performed using Trim Galore! (http://www.bioinformatics.babraham.ac.uk/projects/trim_galore/) script. Gene counts were normalized among various samples using DESeq2 ^40^, and only genes coding for protein and long non-coding RNA (lncRNA) were retained for downstream analysis. Only genes expressed above 1 fragment per kilobase of transcript per million mapped reads (FPKM), in at least one sample, were considered as “expressed” genes and thus included for downstream analysis. Differentially expressed genes (DEGs) were identified using DESeq2, by using as selection parameter adjusted *P*-value lower than 0.01 and likelihood ratio test (LRT) ^40^. Batch effects were removed using the limma package’s “removeBatchEffect” function before performing principal component analysis (PCA). PCA was performed on DEGs by using Bioconductor/R package pcaExplorer v.2.10.0.

### Computational Inference of Developmental Path

Hierarchical clustering was performed using the Euclidean distance and Ward aggregation as criteria. To find a suitable linear order within the hierarchical clustering dendrogram (HSCs, NMs, PMs, MYs, MMs, BCs, SCs and mature blood neutrophils), we used the optimal leaf ordering (OLO) seriation method of R package seriation, version 1.2-9. ^65^. The seriation algorithm is based on the function ‘seriate’ which tries to find a linear order for objects using data in form of a dissimilarity matrix.

### Seven Bridges processing for scRNA-seq data

After demultiplexing of bcl files by using Bcl2fastq2 V2.20 from Illumina and assessment of reads quality, paired-end scRNA-seq reads were then filtered for valid cell barcodes using the barcode whitelist provided by BD. Then, sequenced reads were aligned to the hg38 human transcriptome and the expression of transcripts in each cell was quantified *via* the standard Rhapsody analysis pipeline (BD Biosciences) on Seven Bridges (https://www.sevenbridges.com), following manufacturer’s recommendations.

### Seurat workflow for scRNA-seq data analysis

The R package Seurat ^66^ was utilized for all downstream analysis. For each single cell dataset, the number of detected genes, the number of unique molecular identifiers (UMIs), as well as the fraction of UMIs corresponding to mitochondrial features, which altogether reflect the transcriptome quality of each cell, were calculated. Only cells that transcribed at least 200 genes, and only genes that were expressed in at least 10 cells, were included in the analysis. In sum, 15969 cells and 17398 genes were obtained. Next, cells with more than 25 % mitochondrial transcripts and cells with a number of genes above the 99^th^ percentile or below the 1^st^ percentile were removed for downstream analysis. Therefore, after such a rigorous quality control, a total of 15057 cells were analyzed (4465, 6958 and 3634 from BMs of, respectively, donor 1, donor 2 and donor 3). To examine cell cycle variation in our data, by using CellCycleScoring function we assigned a score to each cell, based on its expression of G2/M and S phase markers. To remove batch effects across data from different donors, we performed dataset integration using SCTransform integration workflow (https://satijalab.org/seurat/v3.2/integration.html). Normalization and detection of high variable genes was performed using the difference between the S and G2M cell cycle score and the percentage of mitochondrial UMIs as covariates. To identify the integration of anchor genes among the 3 datasets from different BMs, the FindIntegrationAnchors() function was used applying default parameters. Using Seurat’s IntegrateData(), samples were combined into one object. These ‘integrated’ batch-corrected values were then set as the ‘default assay’, and gene expression values were scaled before running PCA. The dimensional reduction of the integrated dataset was computed by summarizing the first 50 PCs and visualized in a two-dimensional UMAP representation. Clustering was conducted using the FindNeighbors() and FindClusters() functions using the same 50 PCs, and a resolution parameter set to 0.3. Differential expression (DE) tests were performed using FindAllMarkers() function. DEGs were identified using the non-parametrical Wilcoxon Rank Sum test, based on normalized data. *P* value adjustment was performed using Bonferroni correction bases on the total number of genes in the dataset. Genes with > 0.25 log-fold changes, at least 25% expressed in tested groups, and Bonferroni-corrected p values < 0.01 were considered as significantly DEGs. Euclidean distance between clusters was calculated on DEGs and hierarchical clustering was performed in R, using the ‘ward.D’ linkage clustering. The average gene expression of clusters was calculated using the function AverageExpression().

### Gene ontology (GO) of differentially expressed genes from bulk RNA-seq and scRNA-seq

GO analysis was performed on DEGs from bulk RNA-seq and scRNA-seq for the cellular component, biological process and molecular function ontology domains by using the Bioconductor/R package clusterProfiler (version 3.14.3) ^67^. Over-representation analysis was performed using the Benjamini-Hochberg (BH) procedure, with *P* value Cutoff = 0.01 and Q-value Cutoff = 0.05. Redundant GO terms were removed by using the “simplify” function in the clusterProfiler package, using the Wang similarity measure and a similarity cutoff of 0.5. The most significant terms were then plotted in R.

### Trajectory analysis and Pseudotime calculation

Trajectory analysis was performed by using the destiny algorithm v3.01 ^43^. In brief, the neutrophil progenitor space was subsetted and the diffusion map was calculated based on the top 2000 variable genes with a sum of at least 10 counts over all cells. Based on the diffusion map, a diffusion pseudotime was calculated to infer a transition probability between the different cell states of the neutrophils.

### Statistical analysis

Statistical evaluation was performed by using one-way or two-way analysis of variance, followed by Tukey’s or Bonferroni’s post hoc test, respectively. Values of *P < 0*.*05* were considered statistically significant. Data are expressed as means ± s.e or SEM of the number of indicated experiments. Statistical analysis was performed by Prism Version 7.0 software (GraphPad).

## SUPPLEMENTAL FIGURE LEGENDS

**Figure S1.**
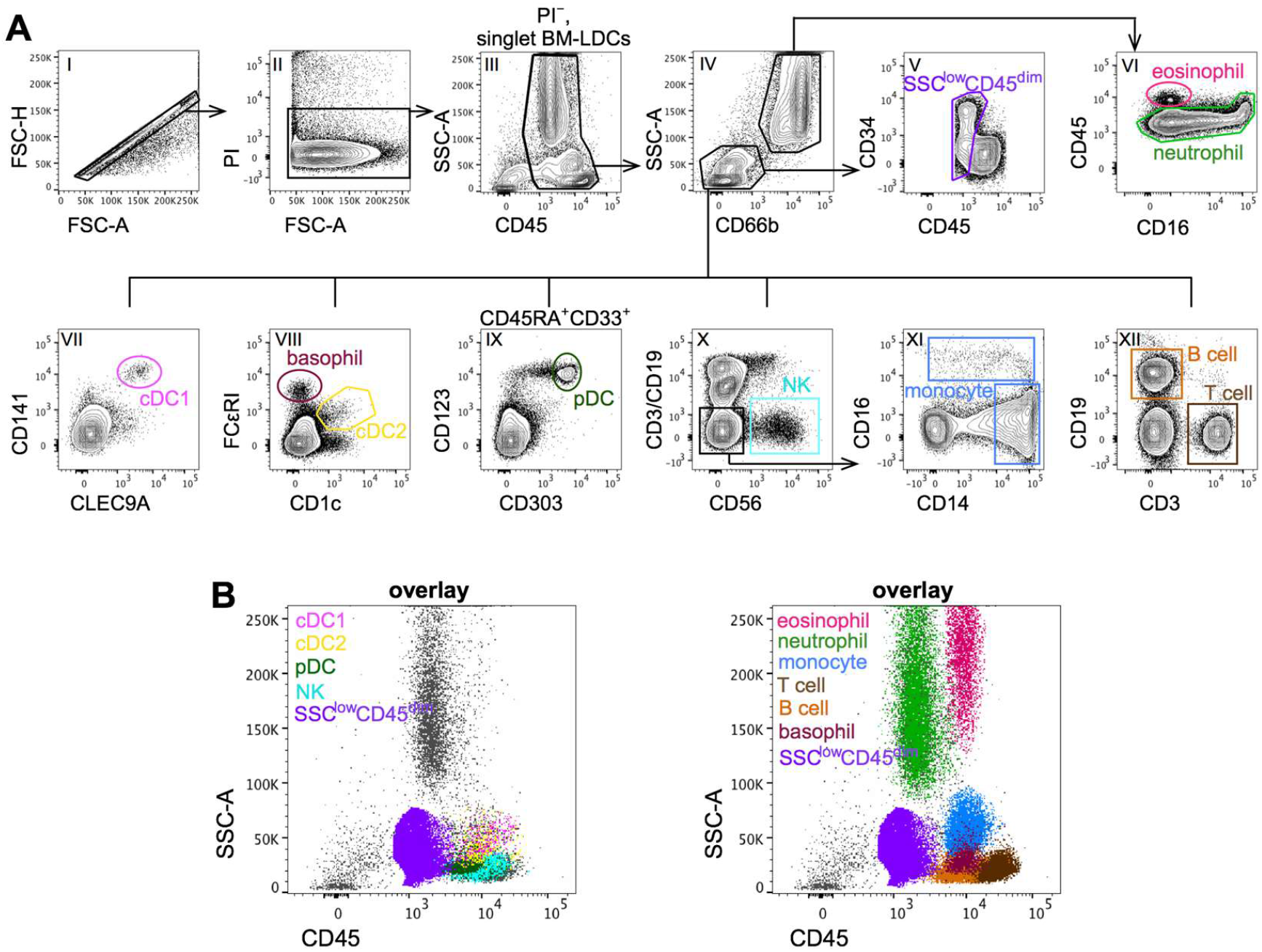
Gating strategy to identify lineage-positive cells and immature SSC^low^CD45^dim^ myeloid/lymphoid progenitors within BM-LDCs. (A) Workflow for the identification of mature leukocyte populations and SSC^low^CD45^dim^ immature myeloid/lymphoid progenitors within the BM. Mature cell populations were identified by sequentially gating on singlets (I), then on PI^−^ cells (II), and finally on CD45^+^ cells (III). Both lineage-positive cells and immature progenitors were identified by gating on either SSC^low^CD66b^−^, or SSC^hi^CD66b^+^, fraction (IV). The sequential steps shown in panels V-XII were done to identify: i) SSC^hi^CD66b^+^CD45^hi^ eosinophils (pink gate in panel VI); ii) SSC^hi^CD66b^+^CD45^+^ neutrophils (green gate in panel VI); iii) SSC^low^CD141^+^CLEC9A^+^ cDC1 DCs (magenta gate in panel VII); iv) SSC^low^FcεRI^+^CD1c^−^ basophils (dark red gate in panel VIII); v) SSC^low^FcεRI^dim^CD1c^+^ cDC2 DCs (yellow gate in panel VIII); vi) SSC^low^CD45RA^+^CD33^+^CD123^+^CD303^+^ pDCs (green gate in panel IX); vii) SSC^low^CD3^−^CD19^−^CD56^+^ NK cells (light blue gate in panel X); viii) SSC^low^CD3^−^CD19^−^ CD56^−^CD14^+^CD16^−^/CD14^dim/−^CD16^+^ total monocytes (blue gates in panel XI); ix) SSC^low^CD3^+^CD19^−^ T cells (brown gate in panel XII); x) SSC^low^CD3^−^CD19^+^ B cells (orange gate in panel XII); and xi) SSC^low^CD45^dim^ cells (purple gate in panel V) including CD34^+^CD38^−^ HSCs, CD34^+^ and CD34^dim/−^ myeloid/lymphoid progenitors. (B) Dot plot overlays of both lineage-positive/mature cells, as determined in (A) (colored populations), and SSC^low^CD45^dim^ immature progenitors (purple population) with total BM-LDCs. Both panels locate lineage-positive cells outside the SSC^low^CD45^dim^ region.

**Figure S2.**
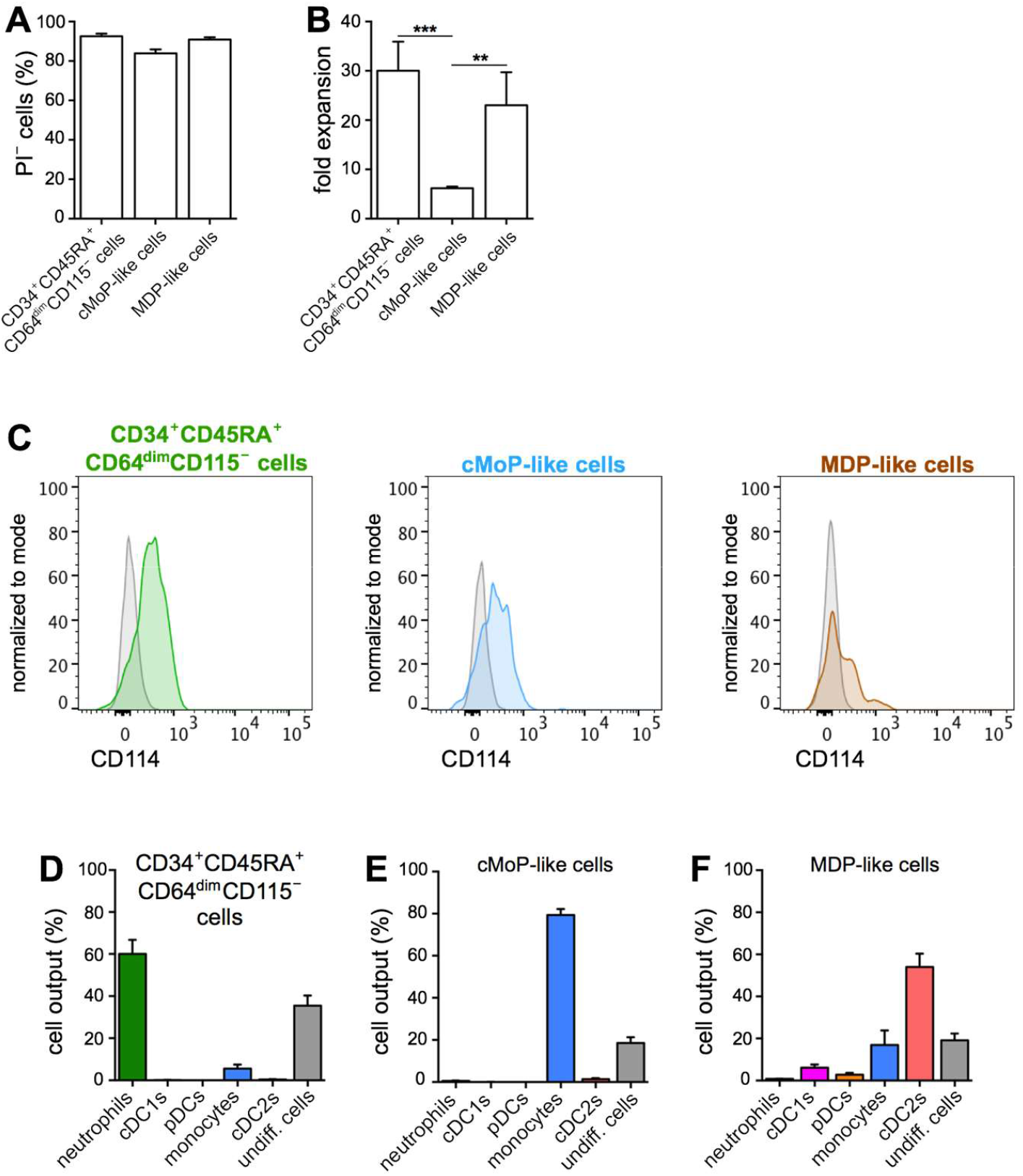
Biological features of CD34^+^CD45RA^+^CD64^dim^CD115^−^, cMoP-like and MDP-like populations, prior to, and after treatment with, either SFGc, or SFG. (A) Bar graphs depict the percentage of viability of the cells generated by a 7 day-treatment of CD34^+^CD45RA^+^CD64^dim^CD115^−^, cMoP-like and MDP-like populations with SFGc (mean ± SEM, relative to CD45^+^ cells, n=6-15). (B) Bar graphs show the expansion of CD34^+^CD45RA^+^CD64^dim^CD115^−^, cMoP-like and MDP-like cells after a 7 day-treatment with SFGc. Values indicate the fold changes of live, CD45^+^ output cells, from the number of input cells (mean ± SEM, n=6-15, **p < 0.01; ***p < 0.001). (C) Histogram overlays showing the expression of CD114 by CD34^+^CD45RA^+^CD64^dim^CD115^−^, cMoP-like and MDP-like populations. Gray profile correspond to isotype control. (D-F) Bar graphs depict the percentage (mean ± SEM, n=7-9, **p < 0.01; ***p < 0.001) of the various mature cell types (listed in the *x* axis) generated after a 7 day-culture with SFG, relative to live, CD45^+^ cells.

**Figure S3:**
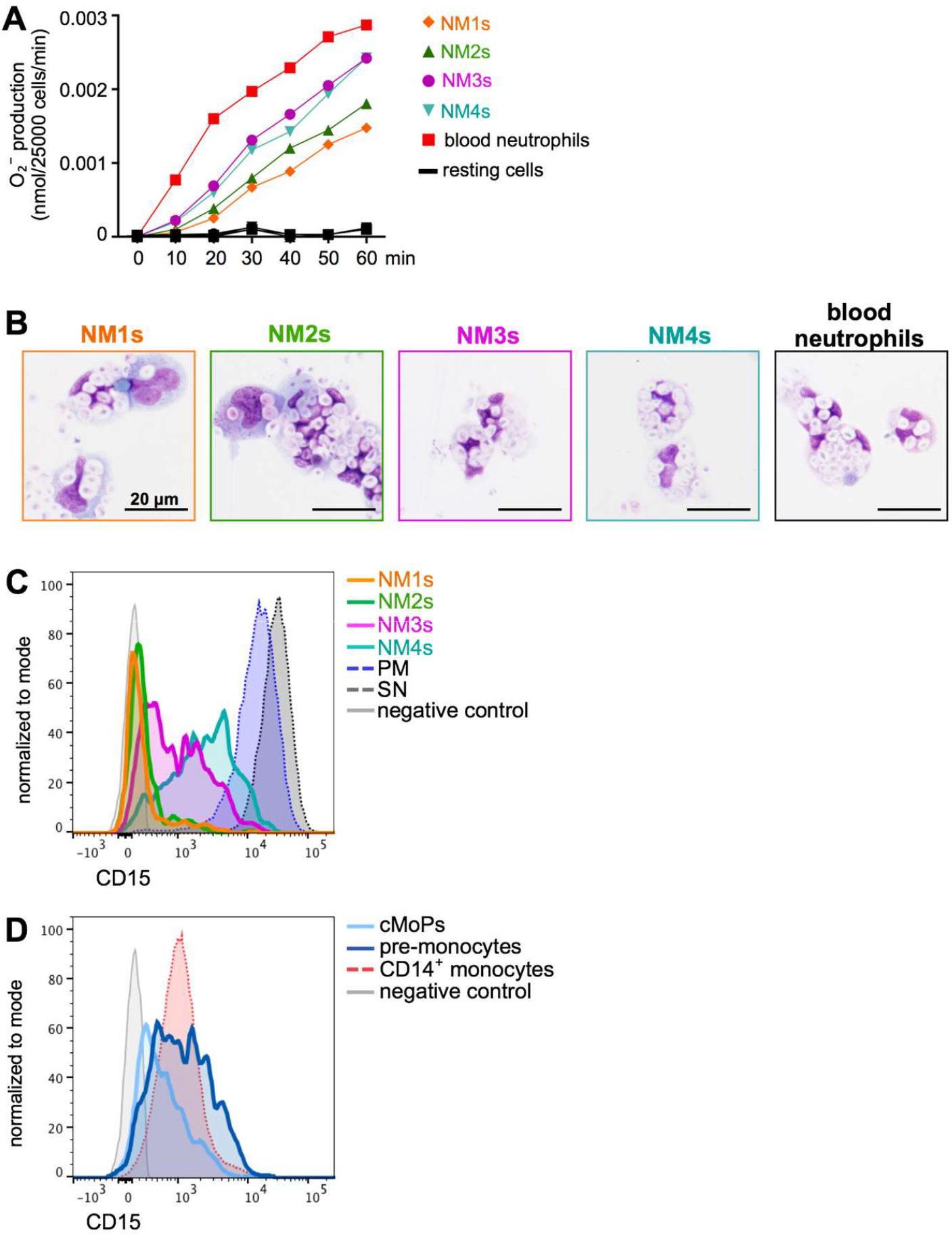
Functionality of neutrophils derived from NMs, as well as CD15 expression by both neutrophil and monocyte progenitors among BM-LDCs and their mature counterparts. (A) Sorted NMs were cultured for 7 days with SFGc. NM-derived neutrophils were recovered as described in M&M, and then stimulated with 20 ng/ml PMA to measure their respiratory burst capacity as compared to that of blood neutrophils freshly purified from HDs. Representative plot (n=3) depicts a time-course of O_2_^−^production. (B) Phagocytic activity of the same NM-derived neutrophils shown in (A), as tested by culturing them with 20 μM unopsonized zymosan particles for 1 h. May-Grunwald Giemsa cytospin preparations show the phagocytosis for each type of NM-derived neutrophils. (C,D) Histogram overlays show the expression of CD15 by the indicated cell populations (represented by the colored lines), as compared to the fluorescence negative control (light gray line). One representative experiment out of 5 with similar results is shown.

**Figure S4:**
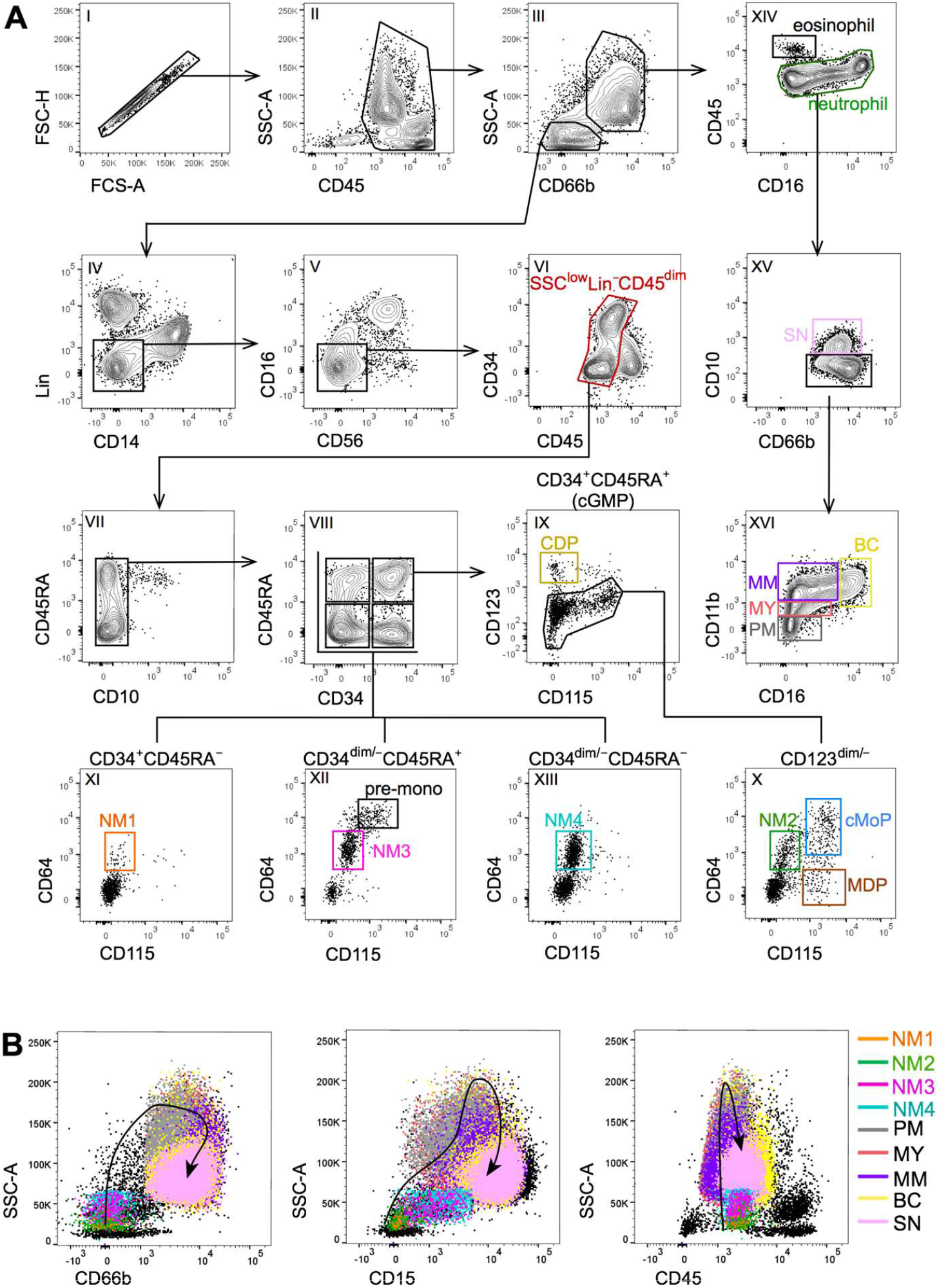
14-color flow cytometry antibody panel to identify NMs, monocyte/DC progenitors and immature neutrophil populations in human BMs. (A) BM-LDCs were stained using a 14-color antibody panel specified in Table S6. Steps shown in panels I-II were sequentially used to exclude doublets (I) and gate on CD45^+^ leukocytes (II). In steps III-V, analysis was performed on SSC^low^CD66b^−^ cells (III), CD3/CD19/CD1c/CD141(Lin)-negative cells (IV) and CD16/CD56-negative cells (V), to exclude lymphoid cells and mature monocytes/DCs. Within the latter cells, the CD45^dim^ region (red gate in panel VI) defines the SSC^low^Lin^−^CD45^dim^ progenitor pool, from which we could subsequently exclude CD45RA^+^CD10^+^ lymphocyte progenitors (VII). Of note, the SSC^low^Lin^−^CD45^dim^ region substantially resembles the equivalent region shown in Figure 1, Figure 2, Figure S1 and Figure S4, thus confirming that both 8-color and 14-color gating strategies are comparable. SSC^low^Lin^−^CD45^dim^CD10^−^ progenitor cells (VII) were then displayed by the CD34/CD45RA marker combination, to separate them in the four region (VIII) equivalent to those shown in Figure 3A. Finally, panels IX-XIII show the strategy used to identify each progenitor population, namely: CDPs (yellow gate in panel IX) within the CD34^+^CD45RA^+^/cGMP fraction; NM2s (green gate in panel X), cMoPs (light blue gate in panel X) and MDPs (brown gate in panel X) within the CD123^dim/−^ fraction of the cGMPs; NM1s within the CD34^+^CD45RA^−^ fraction (orange gate in panel XI); NM3s (magenta gate in panel XII) and pre-monocytes (black gate in panel XII) within the CD34^dim/−^CD45RA^+^ fraction; NM4s within the CD34^dim/−^CD45RA^−^ fraction (cyano gate in panel XIII). We also analyzed the composition of SSC^hi^CD66b^+^ granulocytes (panel III). In steps XIV-XVI, we can identify neutrophil maturation stages from PMs to SNs, by excluding mature eosinophils (XIV). Accordingly, neutrophils (green gate in panel XIV) include CD10^+^ SNs (pink gate in panel XV) and, within the CD10^−^ fraction, PMs (gray gate in panel XVI), MYs (light red in panel XVI), MMs (purple gate in panel XVI) and BCs (yellow gate in panel XVI), all of them identified by analyzing CD11b and CD16 expression. (B) Dot plot overlays depicting all cellular components of the neutrophil maturation cascade, starting from the NMs and up to SNs, as identified in (A). A different color identifies each cell population according to the labels listed on the right of the plot, including total BM-LDCs (depicted in black). Dot plot overlays show the phenotype variation (indicated by the black arrows) of the populations composing the neutrophil maturation cascade, in terms of SSC parameter and CD66b, CD15 and CD45 marker modulation.

**Figure S5:**
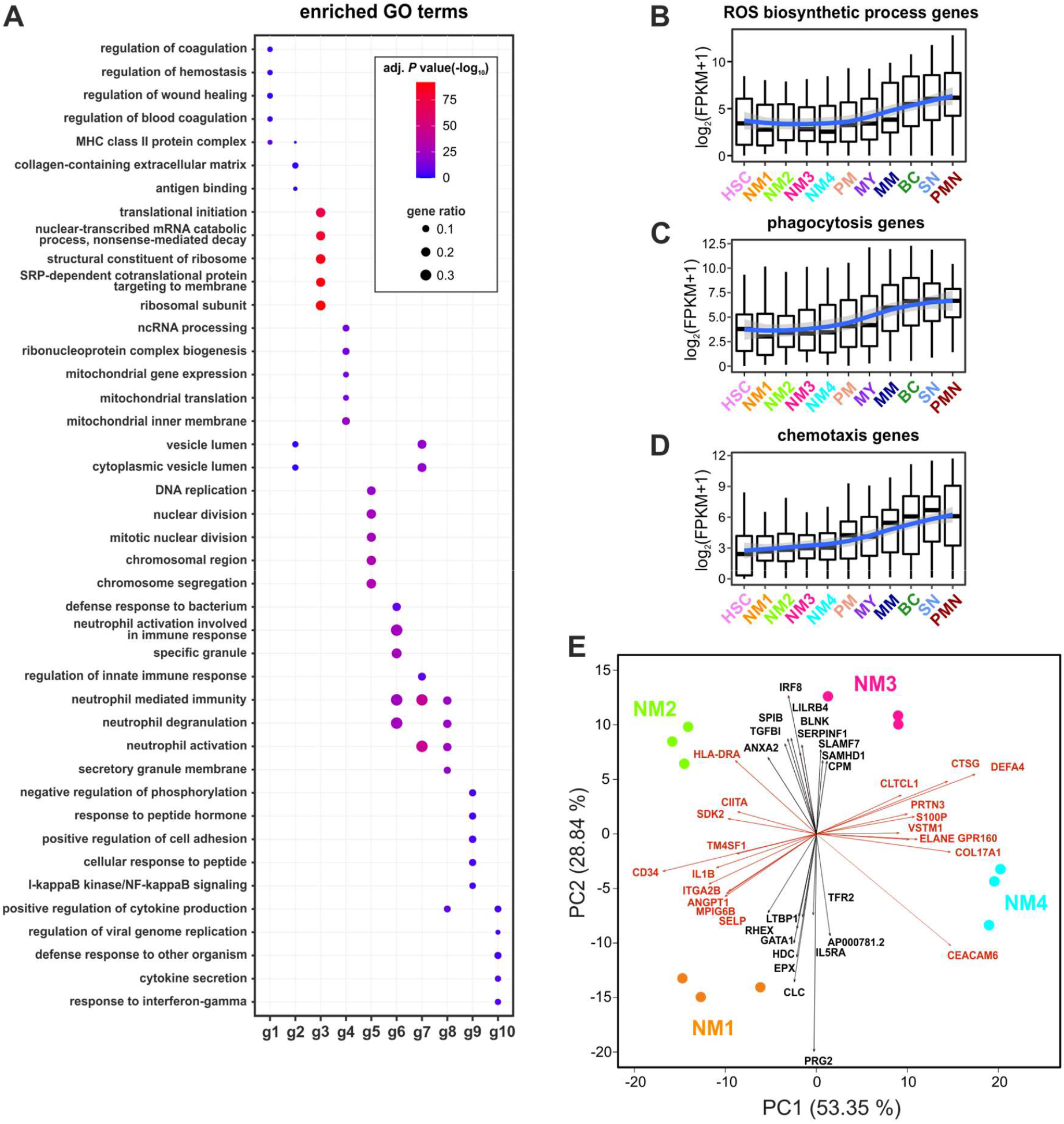
Characterization of NMs transcriptomes as determined by RNA-seq. (A) GO terms enriched by genes associated with the ten gene groups (g1-g10) identified by K-means analysis, as shown in Figure 4C. The top five GO terms with Benjamini-Hochberg-corrected P values <0.05 (one-sided Fisher’s exact test) are shown for every gene group. ‘Gene ratio’ indicate the fraction of DEGs present in the given GO term. (B-D) Box plots showing the distribution of mRNA expression levels [as log2(FPKM+1)] for genes associated to ROS biosynthetic process (B), phagocytosis (C), and chemotaxis (D). Upper and lower boxplot margins indicate first and third quartiles of genes expression levels. LOESS fitting of the data with relative confidence interval is represented by a blue line with a shadow area. (E) PCA biplots based on the DEGs identified among bulk RNA-seq of NM1s, NM2s, NM3s and NM4s. The graph lists the ten most relevant genes contributing to sample variations (indicated by vectors) for both PC1 and PC2, under both positive and negative directions. Vector lengths correlate with the weight of the given gene within the components.

**Figure S6:**
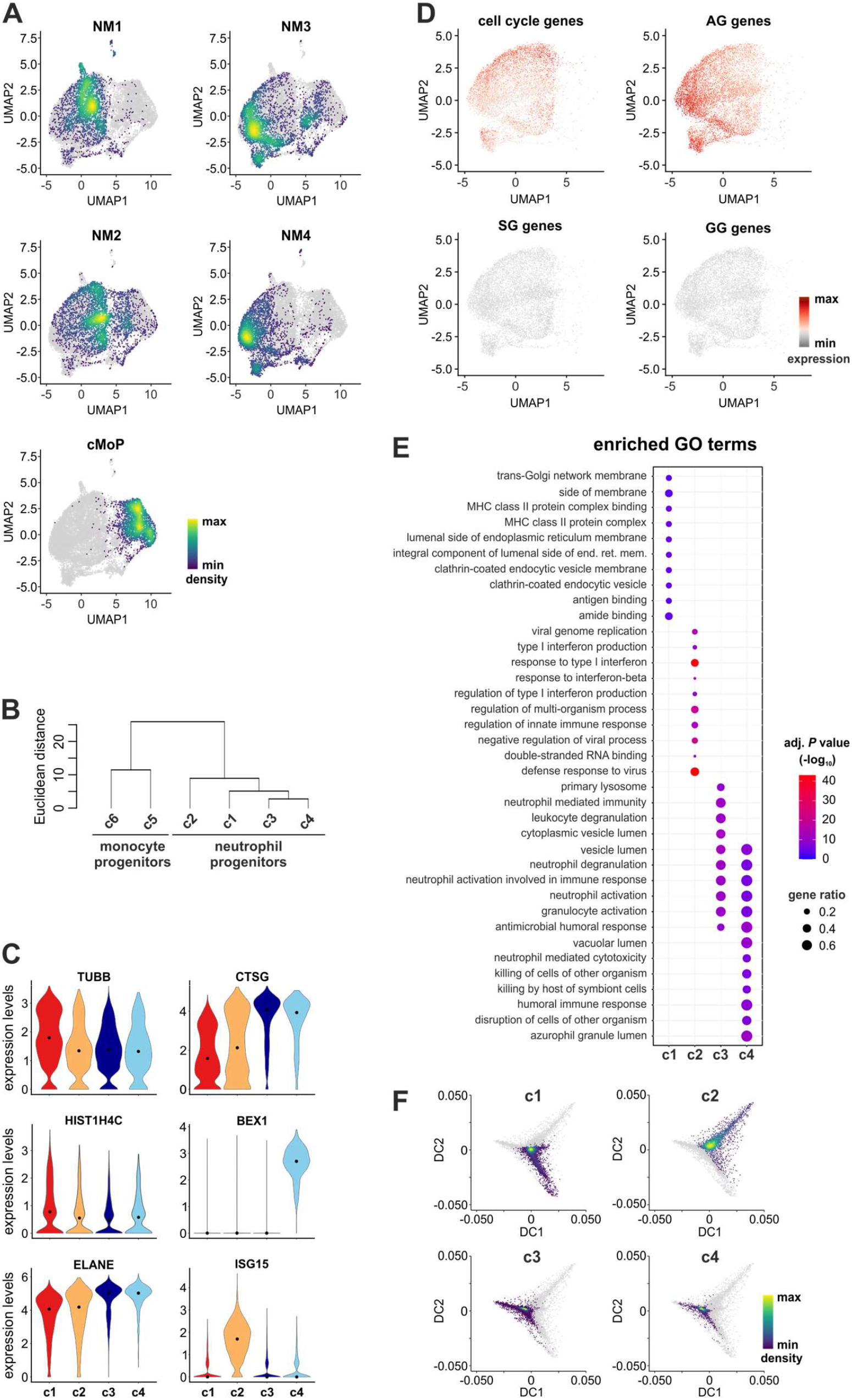
Additional characterization of NM and cMoP scRNA-seqs. (A) Density plots of NM1s, NM2s, NM3s, NM4s and cMoPs overlaid on the UMAP of Figure 5A. Density of cells in each plot is depicted according to the indications of the color bar. (B) Hierarchical clustering dendrogram based on the DEGs identified among the neutrophilic and monocytic cell clusters shown in Figure 5B. The vertical axis of the dendrogram represents the dissimilarity between clusters (i.e., Euclidean distance) (C) Violin plots showing the expression of selected genes across the four neutrophil clusters (c1-c4) chosen among the top defining genes indicated in Figure 6E. (D) Expression patterns of cell-cycle, AG, SG, and GG genes projected on the UMAP plot restricted to neutrophil progenitor clusters (c1-c4) (E) Gene Ontology analysis of DEGs for c2-c4. For every cluster (x-axis), the top ten Gene Ontology terms with Benjamini-Hochberg-corrected P values <0.05 (one-sided Fisher’s exact test) are shown. For cluster 1 no enrichment of biological processes GO term was identified (F) Trajectory plots of c1-c4 cells as defined in Figure 5B. In each plot is depicted the density of cells according to the color bar placed at the right bottom corner of the panel

## REFERENCES

1. Notta F, Zandi S, Takayama N, et al. Distinct routes of lineage development reshape the human blood hierarchy across ontogeny. Science. 2016;351:aab2116.

2. Paul F, Arkin Y, Giladi A, et al. Transcriptional Heterogeneity and Lineage Commitment in Myeloid Progenitors. Cell. 2015;163:1663–77.

3. Velten L, Haas SF, Raffel S, et al. Human haematopoietic stem cell lineage commitment is a continuous process. Nat Cell Biol. 2017;19:271–281.

4. Pellin D, Loperfido M, Baricordi C, et al. A comprehensive single cell transcriptional landscape of human hematopoietic progenitors. Nat Commun. 2019;10:2395.

5. Yanez A, Coetzee SG, Olsson A, et al. Granulocyte-Monocyte Progenitors and Monocyte-Dendritic Cell Progenitors Independently Produce Functionally Distinct Monocytes. Immunity. 2017;47:890–902 e4.

6. Zheng S, Papalexi E, Butler A, Stephenson W, Satija R. Molecular transitions in early progenitors during human cord blood hematopoiesis. Mol Syst Biol. 2018;14:e8041.

7. Karamitros D, Stoilova B, Aboukhalil Z, et al. Single-cell analysis reveals the continuum of human lympho-myeloid progenitor cells. Nat Immunol. 2018;19:85–97.

8. Weinreb C, Rodriguez-Fraticelli A, Camargo FD, Klein AM. Lineage tracing on transcriptional landscapes links state to fate during differentiation. Science. 2020;367.

9. Muench DE, Olsson A, Ferchen K, et al. Mouse models of neutropenia reveal progenitor-stage-specific defects. Nature. 2020;582:109–114.

10. Doulatov S, Notta F, Laurenti E, Dick JE. Hematopoiesis: a human perspective. Cell Stem Cell. 2012;10:120–36.

11. Watcham S, Kucinski I, Gottgens B. New insights into hematopoietic differentiation landscapes from single-cell RNA sequencing. Blood. 2019;133:1415–1426.

12. Laurenti E and Gottgens B. From haematopoietic stem cells to complex differentiation landscapes. Nature. 2018;553:418–426.

13. Jacobsen SEW and Nerlov C. Haematopoiesis in the era of advanced single-cell technologies. Nat Cell Biol. 2019;21:2–8.

14. Manz MG, Miyamoto T, Akashi K, Weissman IL. Prospective isolation of human clonogenic common myeloid progenitors. Proc Natl Acad Sci U S A. 2002;99:11872–7.

15. Doulatov S, Notta F, Eppert K, et al. Revised map of the human progenitor hierarchy shows the origin of macrophages and dendritic cells in early lymphoid development. Nat Immunol. 2010;11:585–93.

16. Lee J, Breton G, Oliveira TY, et al. Restricted dendritic cell and monocyte progenitors in human cord blood and bone marrow. J Exp Med. 2015;212:385–99.

17. Kawamura S, Onai N, Miya F, et al. Identification of a Human Clonogenic Progenitor with Strict Monocyte Differentiation Potential: A Counterpart of Mouse cMoPs. Immunity. 2017;46:835–848 e4.

18. Mori Y, Iwasaki H, Kohno K, et al. Identification of the human eosinophil lineage-committed progenitor: revision of phenotypic definition of the human common myeloid progenitor. J Exp Med. 2009;206:183–93.

19. Gorgens A, Radtke S, Mollmann M, et al. Revision of the human hematopoietic tree: granulocyte subtypes derive from distinct hematopoietic lineages. Cell Rep. 2013;3:1539–52.

20. Drissen R, Thongjuea S, Theilgaard-Monch K, Nerlov C. Identification of two distinct pathways of human myelopoiesis. Sci Immunol. 2019;4.

21. Olweus J, Thompson PA, Lund-Johansen F. Granulocytic and monocytic differentiation of CD34hi cells is associated with distinct changes in the expression of the PU.1-regulated molecules, CD64 and macrophage colony-stimulating factor receptor. Blood. 1996;88:3741–54.

22. Evrard M, Kwok IWH, Chong SZ, et al. Developmental Analysis of Bone Marrow Neutrophils Reveals Populations Specialized in Expansion, Trafficking, and Effector Functions. Immunity. 2018;48:364–379 e8.

23. Zhu YP, Padgett L, Dinh HQ, et al. Identification of an Early Unipotent Neutrophil Progenitor with Pro-tumoral Activity in Mouse and Human Bone Marrow. Cell Rep. 2018;24:2329–2341 e8.

24. Kwok I, Becht E, Xia Y, et al. Combinatorial Single-Cell Analyses of Granulocyte-Monocyte Progenitor Heterogeneity Reveals an Early Uni-potent Neutrophil Progenitor. Immunity. 2020;53:303–318 e5.

25. Dinh HQ, Eggert T, Meyer MA, et al. Coexpression of CD71 and CD117 Identifies an Early Unipotent Neutrophil Progenitor Population in Human Bone Marrow. Immunity. 2020;53:319–334 e6.

26. van Lochem EG, van der Velden VH, Wind HK, et al. Immunophenotypic differentiation patterns of normal hematopoiesis in human bone marrow: reference patterns for age-related changes and disease-induced shifts. Cytometry B Clin Cytom. 2004;60:1–13.

27. Gorczyca W, Sun ZY, Cronin W, et al. Immunophenotypic pattern of myeloid populations by flow cytometry analysis. Methods Cell Biol. 2011;103:221–66.

28. Olweus J, BitMansour A, Warnke R, et al. Dendritic cell ontogeny: a human dendritic cell lineage of myeloid origin. Proc Natl Acad Sci U S A. 1997;94:12551–6.

29. Breton G, Lee J, Liu K, Nussenzweig MC. Defining human dendritic cell progenitors by multiparametric flow cytometry. Nat Protoc. 2015;10:1407–22.

30. Lee J, Zhou YJ, Ma W, et al. Lineage specification of human dendritic cells is marked by IRF8 expression in hematopoietic stem cells and multipotent progenitors. Nat Immunol. 2017;18:877–888.

31. Mora-Jensen H, Jendholm J, Fossum A, et al. Technical advance: immunophenotypical characterization of human neutrophil differentiation. J Leukoc Biol. 2011;90:629–34.

32. Bonifacio M, Stagno F, Scaffidi L, Krampera M, Di Raimondo F. Management of Chronic Myeloid Leukemia in Advanced Phase. Front Oncol. 2019;9:1132.

33. Jamieson CH, Ailles LE, Dylla SJ, et al. Granulocyte-macrophage progenitors as candidate leukemic stem cells in blast-crisis CML. N Engl J Med. 2004;351:657–67.

34. Kinstrie R, Karamitros D, Goardon N, et al. Heterogeneous leukemia stem cells in myeloid blast phase chronic myeloid leukemia. Blood Adv. 2016;1:160–169.

35. Diaz-Blanco E, Bruns I, Neumann F, et al. Molecular signature of CD34(+) hematopoietic stem and progenitor cells of patients with CML in chronic phase. Leukemia. 2007;21:494–504.

36. Bar-Joseph Z, Gifford DK, Jaakkola TS. Fast optimal leaf ordering for hierarchical clustering. Bioinformatics. 2001;17 Suppl 1:S22–9.

37. Olweus J, Lund-Johansen F, Terstappen LW. Expression of cell surface markers during differentiation of CD34^+^, CD38-/lo fetal and adult bone marrow cells. Immunomethods. 1994;5:179–88.

38. Cowland JB and Borregaard N. Granulopoiesis and granules of human neutrophils. Immunol Rev. 2016;273:11–28.

39. Cassatella MA, Ostberg NK, Tamassia N, Soehnlein O. Biological Roles of Neutrophil-Derived Granule Proteins and Cytokines. Trends Immunol. 2019;40:648–664.

40. Love MI, Huber W, Anders S. Moderated estimation of fold change and dispersion for RNA-seq data with DESeq2. Genome Biol. 2014;15:550.

41. Becht E, McInnes L, Healy J, et al. Dimensionality reduction for visualizing single-cell data using UMAP. Nat Biotechnol. 2018.

42. Hafemeister C and Satija R. Normalization and variance stabilization of single-cell RNA-seq data using regularized negative binomial regression. Genome Biol. 2019;20:296.

43. Angerer P, Haghverdi L, Buttner M, et al. destiny: diffusion maps for large-scale single-cell data in R. Bioinformatics. 2016;32:1241–3.

44. Ng LG, Ostuni R, Hidalgo A. Heterogeneity of neutrophils. Nat Rev Immunol. 2019;19:255–265.

45. Goardon N, Marchi E, Atzberger A, et al. Coexistence of LMPP-like and GMP-like leukemia stem cells in acute myeloid leukemia. Cancer Cell. 2011;19:138–52.

46. Schulte-Schrepping J, Reusch N, Paclik D, et al. Severe COVID-19 Is Marked by a Dysregulated Myeloid Cell Compartment. Cell. 2020;182:1419–1440 e23.

47. Lindblad O, Li T, Su X, et al. BEX1 acts as a tumor suppressor in acute myeloid leukemia. Oncotarget. 2015;6:21395–405.

48. Xiao Q, Hu Y, Liu Y, et al. BEX1 promotes imatinib-induced apoptosis by binding to and antagonizing BCL-2. PLoS One. 2014;9:e91782.

49. Xie X, Shi Q, Wu P, et al. Single-cell transcriptome profiling reveals neutrophil heterogeneity in homeostasis and infection. Nat Immunol. 2020;21:1119–1133.

50. Gupta S, Nakabo S, Blanco LP, et al. Sex differences in neutrophil biology modulate response to type I interferons and immunometabolism. Proc Natl Acad Sci U S A. 2020;117:16481–16491.

51. Zilionis R, Engblom C, Pfirschke C, et al. Single-Cell Transcriptomics of Human and Mouse Lung Cancers Reveals Conserved Myeloid Populations across Individuals and Species. Immunity. 2019;50:1317–1334 e10.

52. Mistry P, Nakabo S, O’Neil L, et al. Transcriptomic, epigenetic, and functional analyses implicate neutrophil diversity in the pathogenesis of systemic lupus erythematosus. Proc Natl Acad Sci U S A. 2019;116:25222–25228.

53. Giustacchini A, Thongjuea S, Barkas N, et al. Single-cell transcriptomics uncovers distinct molecular signatures of stem cells in chronic myeloid leukemia. Nat Med. 2017;23:692–702.

54. Calzetti F, Tamassia N, Arruda-Silva F, Gasperini S, Cassatella MA. The importance of being “pure” neutrophils. J Allergy Clin Immunol. 2017;139:352–355 e6.

55. Hochhaus A, Baccarani M, Silver RT, et al. European LeukemiaNet 2020 recommendations for treating chronic myeloid leukemia. Leukemia. 2020;34:966–984.

56. Arber DA, Orazi A, Hasserjian R, et al. The 2016 revision to the World Health Organization classification of myeloid neoplasms and acute leukemia. Blood. 2016;127:2391–405.

57. Zimmermann M, Arruda-Silva F, Bianchetto-Aguilera F, et al. IFNalpha enhances the production of IL-6 by human neutrophils activated via TLR8. Sci Rep. 2016;6:19674.

58. Tura O, Barclay GR, Roddie H, Davies J, Turner ML. Optimal ex vivo expansion of neutrophils from PBSC CD34^+^ cells by a combination of SCF, Flt3-L and G-CSF and its inhibition by further addition of TPO. J Transl Med. 2007;5:53.

59. Jie Z, Zhang Y, Wang C, et al. Large-scale ex vivo generation of human neutrophils from cord blood CD34^+^ cells. PLoS One. 2017;12:e0180832.

60. Serra MC, Calzetti F, Ceska M, Cassatella MA. Effect of substance P on superoxide anion and IL-8 production by human PMNL. Immunology. 1994;82:63–9.

61. Tamassia N, Le Moigne V, Calzetti F, et al. The MyD88-independent pathway is not mobilized in human neutrophils stimulated via TLR4. J Immunol. 2007;178:7344–56.

62. Arruda-Silva F, Bianchetto-Aguilera F, Gasperini S, et al. Human Neutrophils Produce CCL23 in Response to Various TLR-Agonists and TNFalpha. Front Cell Infect Microbiol. 2017;7:176.

63. Picelli S, Faridani OR, Bjorklund AK, et al. Full-length RNA-seq from single cells using Smart-seq2. Nat Protoc. 2014;9:171–81.

64. Bianchetto-Aguilera F, Tamassia N, Gasperini S, et al. Deciphering the fate of slan(+) - monocytes in human tonsils by gene expression profiling. FASEB J. 2020;34:9269–9284.

65. Hahsler M, Hornik K, Buchta C. Getting things in order: An introduction to the R package seriation. Journal of Statistical Software. 2008;25:1–34.

66. Butler A, Hoffman P, Smibert P, Papalexi E, Satija R. Integrating single-cell transcriptomic data across different conditions, technologies, and species. Nat Biotechnol. 2018;36:411–420.

67. Yu G, Wang LG, Han Y, He QY. clusterProfiler: an R package for comparing biological themes among gene clusters. OMICS. 2012;16:284–7.

